# Lipid metabolism of hepatocyte-like cells supports intestinal tumor growth in *Drosophila*

**DOI:** 10.1101/2025.04.04.647255

**Authors:** K. Huang, T. Miao, Y. Chen, E. Dantas, J. Sanford, John M. Asara, SJ. Moon, Y. Hu, K. Wang, M. Han, M. Goncalves, N Perrimon

## Abstract

Tumors reprogram lipid metabolism in distant tissues to support their growth. In adult *Drosophila*, gut tumors secrete the PDGF/VEGF-like factor Pvf1, which activates the TORC1-Hnf4 pathway in hepatocyte-like oenocytes. This drives production of very long-chain fatty acids and wax esters essential for tracheal growth around the tumor. Blocking Hnf4 or the elongase mElo in oenocytes strongly suppresses tracheogenesis, tumor progression, and cachexia-like organ wasting, while extending host lifespan. The same pathway also controls tracheal development in healthy flies. Lipoprotein receptor *LpR2* depletion in oenocytes rescues the observed tumor induced tracheal tracheal remodeling. This tumor-host interaction is conserved: VEGF-A induces lipid metabolism genes in human hepatocytes, and lung tumor-bearing mice show elevated hepatic *Hnf4* and *Elovl7*. The study reveals a non-autonomous role of the TORC1-Hnf4 axis in lipid-mediated tumor progression and identifies potential therapeutic targets for cancer-associated metabolic dysfunction.

## Introduction

Tumor growth demands a growing supply of nutrients and oxygen. The growing tumor constantly shapes its microenvironment to favor fast growth and expansion, through secreted factors and extracellular matrix [1–4]. The composition of the tumor microenvironment (TME) varies between tumor types, but the hallmarks include increased blood vessels, immune cells, stromal cells, and extracellular matrix [5]. To overcome the increasingly hypoxic and acidic environment, TME promotes angiogenesis to restore oxygen and nutrient supply. In addition to modulating the local TME, malignant cells can exert systemic effects on distant organs—including the liver, skeletal muscle, and brain—to reshape the host’s metabolic pathways, ultimately promoting further tumor progression [6–10]. Liver is a metabolic center for amino acids, carbohydrates, and lipid. A key liver-specific metabolic function is the complete urea cycle (UC), which eliminates excess ammonia by converting into urea, a less toxic waste. Previous studies have identified evidence of decreased UC activity in the livers of 4T1 breast cancer bearing mice, and the plasma of children with cancer [9], demonstrating an existing interaction between tumors and the liver [11–15]. Delineating molecular pathways involving tumor-host interaction can provide therapeutic targets.

Lipids play a critical role in cancer progression, by serving as building blocks, energy sources and signaling molecules. It has been observed that tumor cells reprogram lipid metabolism to enhance growth, characterized by increased lipid synthesis, lipid uptake, fatty acid oxidation (FAO) and lipid storage [16]. Tumor cells also actively reshape lipid metabolism in cells from the TME through secreted signaling molecules and drive TME to favor tumor growth. For example, increased lipid uptake and FAO in regulatory T cells (Tregs), tumor-associated macrophages (TAMs) and myeloid-derived suppressor cells (MDSCs), promote their immunosuppressive function [17–19]. However, whether tumor cells also reshape lipid metabolism in distal organs, such as liver, which is a metabolic center for lipid transport, is not known.

The adult *Drosophila* midgut is a powerful model for investigating tumorigenesis and tumor-driven metabolic remodeling. Many genetic alterations associated with human cancers induce hyperproliferation of fly intestinal stem cells (ISCs) and lead to tumor formation in the midgut [20]. In particular, expression of constitutively active Yorkie (yki^3SA^)—the *Drosophila* homolog of human YAP1—drives gut tumor development and triggers cachexia-like phenotypes, including lipid depletion, hyperglycemia, and wasting of muscle and ovaries [21, 22]. Mechanistically, yki^3SA^ gut tumors secrete multiple systemic factors, such as the IGF-antagonizing peptide ImpL2 [21], PDGF- and VEGF-related factor 1 (Pvf1) [22] and unpaired 3 (Upd3), a functional ortholog of mammalian interleukin-6 (IL-6) [22]. Notably, IL-6- and PDGF/VEGF-associated inflammatory programs are also broadly elevated across diverse cancer models [23–26]. Together, these observations highlight tumor–host organ communication as an evolutionarily conserved process and establish *Drosophila* as an excellent system to dissect tumor–host interactions *in vivo*.

As *Drosophila* has emerged as a simple yet powerful model to study tumor-host interactions [27], we use this model to better understand the reciprocal communication between liver and tumor cells. The fly midgut, oenocytes, fat body and Malpighian tubules are functionally equivalent to human digestive tract, hepatocytes, adipose tissue and kidney, respectively [28]. Oenocytes, which are hepatocyte-like cells, are important for mobilizing stored lipids during starvation and interact closely with metabolic tissues, such as the fat body to regulate lipid uptake and reprocessing [29, 30]. In addition to regulating lipid metabolism under nutritional deprivation, oenocytes are the major site for synthesis of cuticular hydrocarbons in pterygote insects [31–33]. Adult oenocytes synthesize hydrocarbons using fatty acyl-CoA precursors via reduction to fatty aldehyde and oxidative decarbonylation [34, 35]. Many fatty acid synthases, elongases and desaturases expressed in adult oenocytes produce various types of fatty acyl-CoA substrates that are processed to form a complex blend of cuticular hydrocarbons [36–39]. Hepatocyte nuclear factor 4 (Hnf4), an oenocyte enriched transcription factor, plays an important role in regulating the conversion of lipids to very long chain fatty acid, which are important substrates for hydrocarbon synthesis [39]. Oenocytes are the major site for lipid transport and processing, reminiscent of human hepatocytes. However, whether and how oenocytes regulate tumor growth is unknown.

The *Drosophila* trachea is a network of oxygen transporting tubes, the functional equivalent of mammalian blood vessels [40]. Trachea is composed of terminal tracheal cells (TTCs), which are highly plastic cells, and the terminal branches (TBs) or tracheoles which deliver oxygen to the tissues. TTCs are oxygen sensing cells, and hypoxic conditions promote the formation of TBs through hypoxia-inducible factor-1α (HIF-1α; Similar (Sima) in *Drosophila*), similar to the role of HIF-1α in mammalian angiogenesis. Trachea resides in proximity of many tissues, including the midgut. Trachea penetrate the visceral muscles to access the intestinal epithelium to supply it with oxygen [41]. The intestinal epithelium is apicobasally polarized, composed of ISCs and their progeny. Damage induced by enteric infection, oxidative agents and tumor growth can increase tracheogenesis on the midgut, which is necessary for efficient damage-induced ISC-mediated regeneration [42, 43].

Additionally, tumor-induced tracheogenesis in *Drosophila* is reminiscent of cancer-induced angiogenesis in mammals [42].

Using a *Drosophila* gut tumor model, we uncover a mechanism through which gut tumors reprogram lipid metabolism in distal oenocytes to promote both tracheal development and tumor growth. We find that tumors secrete a PDGF/VEGF-like factor, Pvf1, which activates the TORC1-Hnf4 signaling pathway in oenocytes. This activation boosts the production of specific lipids, including very long-chain fatty acids and wax esters, essential for tracheal growth around the tumor. Notably, reducing the activity of Hnf4 and mElo, an elongase involved in generating very long-chain fatty acids, in oenocytes suppress tumor growth, tracheal development, and cachexia-like phenotypes, while also extending lifespan. Additionally, we demonstrate that this regulatory pathway is conserved in mammals, as VEGF-A stimulates lipid metabolism gene expression in human hepatocytes, and tumor-bearing mice exhibit increased hepatic expression of lipid elongation genes. Our findings highlight a tumor-host interaction, where tumors non-autonomously reprogram distal lipid metabolism to facilitate their growth.

## Results

### Oenocyte lipid metabolism regulated by Hnf4 controls tumor proliferation

To investigate how tumor growth rewires lipid metabolism of the whole fly, we expressed an active form of *Yorkie ( yki^3SA^),* in adult gut stem cells, using the intestinal stem cells (ISCs)/enteroblast driver (*Esg-Gal4* or *Esg-LexA*), hereafter referred to as Yki flies [21]. *Esg-Gal4* or *Esg-LexA* was combined with the *tubulin* promoter-driven temperature-sensitive *GAL80 (tubP-GAL80^TS^)* to control the temporal expression of Gal4 or LexA. Next, we conducted untargeted lipidomic profiling using an LC-MS/MS-based platform of whole-body samples of adult female flies (*Esg-LexA > LexAop-yki^3SA^*) expressing *yki^3SA^* for 3 days, together with control flies (*Esg > +*). The complete *Drosophila* genotype information can be found in Supplementary Data d 1. Female flies are used in the analysis unless otherwise noted because they show more significant and consistent tumor-associated cachexia phenotypes. We detected 2471 lipid species in total representing 45 lipid classes (Supplementary Data 2). The lipidomic analysis showed an increase in wax ester (WE), monogalactosyldiacylglycerol (MGDG), phosphatidylglycerol (PG), phosphatidylcholine (PC), ceramide (Cer) and acylcarnitine (AcCa) levels in Yki flies (Figure 1A). We also observed a slight reduction of triglycerides (TG) in Yki flies at early stage, in agreement with previous findings [21, 44]. WEs are major components of cuticular lipids, which cover nearly all parts of terrestrial arthropods [45] to restrict water loss, and WE levels are significantly increased in Yki flies (Figure 1B), yet the significance of this regulation is unknown. By profiling the fatty acid carbon chain length of WEs, we found that Yki flies exhibit elevated levels of WEs containing very-long-chain fatty acids (carbon length ≥22), as well as WEs with shorter chains (<22) (Figure 1C). As the biosynthesis of cuticular lipids mainly occurs in *Drosophila* oenocytes and Hnf4 is a main regulator of this process [39], we tested whether oenocyte Hnf4 regulates the level of WEs in control flies. WEs were slightly reduced when *Hnf4* was knocked down in control adult oenocytes using the *PromE(800)-Gal4* temperature sensitive driver (*PromE-GAL4,Tub-GAL80^TS^*, referred to as *PromE*). PromE(800)-Gal4 is specifically expressed in adult female oenocytes, although it is also expressed in the reproductive organs of adult males [31]. To ensure tissue specificity, female flies were used in all experiments unless otherwise noted. Next, to test whether Hnf4 regulates WE synthesis in Yki flies, we used the LexA-LexAop system to induce Yki tumors in the gut and the Gal4/UAS system in oenocytes (*PromE-GAL4, Tub-GAL80^TS^*, referred to as *PromE*) to knockdown *Hnf4*. Knockdown of *Hnf4* in oenocytes decreased Yki-induced WE levels (Figure 1D), indicating that *Hnf4* expression in oenocytes is required for the increase of WE.

**Figure 1:**
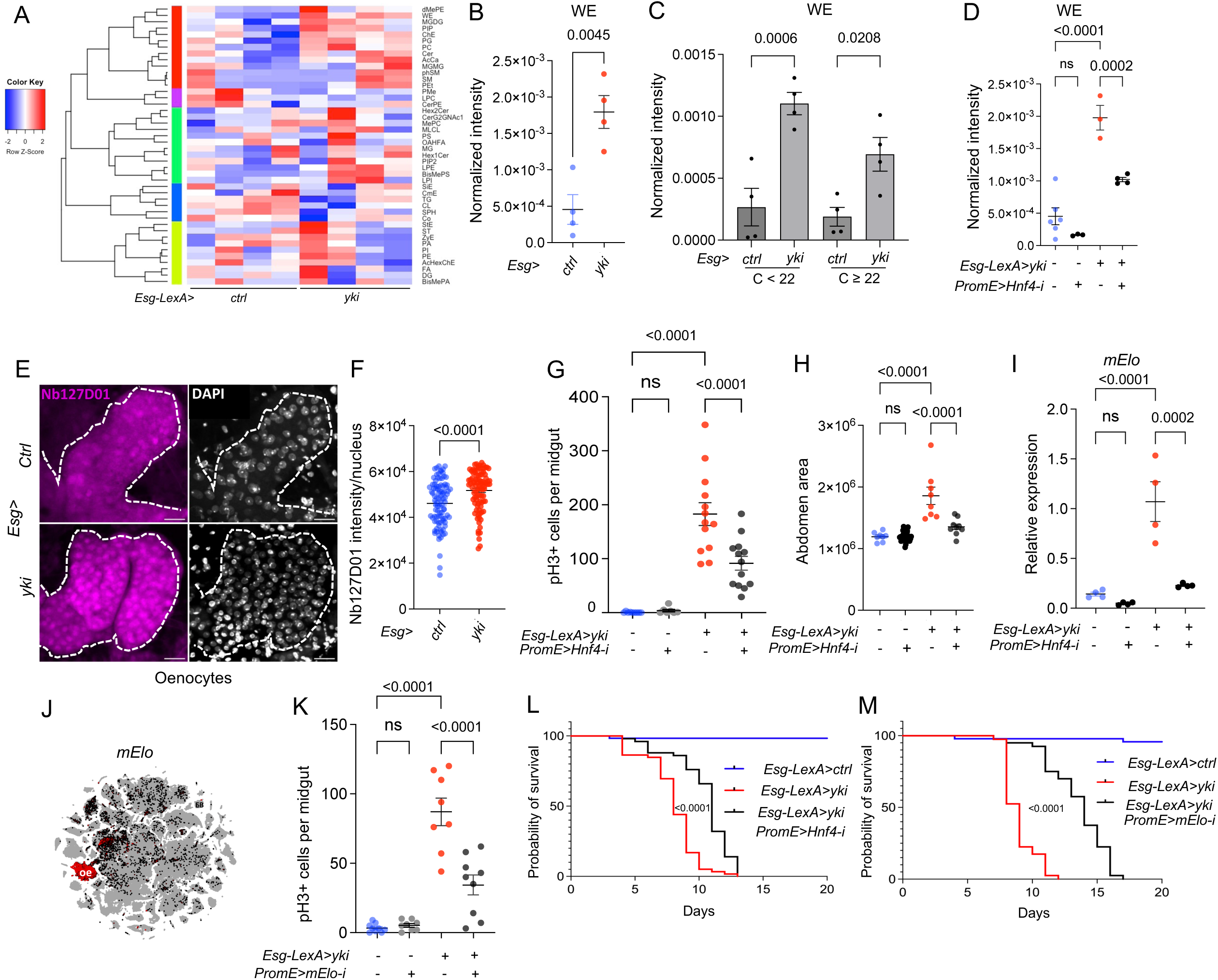
Oenocyte Hnf4 regulates wax ester synthesis and supports Yki-induced tumor growth. (A-B) Heatmap of the normalized total ion intensity of major lipid classes identified in whole bodies from control flies (*Esg-LexA*) or Yki flies (*Esg-LexA/LexAop-yki^S3A^; PromE-GAL4, Tub-GAL80^TS^/+*), with four biological replicates per condition. Scaled colors are presented as the values relative to the average of each row. For the full names corresponding to the abbreviations, see Supplementary Data 4. (B) Untargeted lipidomic profiling using LC-MS/MS of whole-body extracts revealed increased wax ester (WE) levels in Yki flies. Normalized intensity was calculated as the integrated LC–MS peak area for each lipid species, normalized to the sample’s total signal (total reads) and to the analyzed sample volume. P = 0.0045. (C) Fatty acid chain-length distribution (C< 22 and C≥22) in detected WE level from control and Yki flies. Adjusted P values from left to right: 0.0006, 0.0208. (D) WE level in control flies, Yki flies, and *Hnf4* knockdown in oenocytes of Yki flies, N = 4 biological replicates. Adjusted P values from left to right: <0.0001, 0.0002. (E-F) Immunostaining of Nb127D01 for nuclear-localized, endogenously tagged Hnf4 (Hnf4-127D01) in oenocytes in controls and in Yki flies. DAPI staining marks nuclei, with dashed lines enclosing oenocytes. N = 8 independent experiments. P < 0.0001. (G) Quantification of intestinal stem cell (ISC) proliferation, as indicated by phospho-histone H3 (pH3+) cells in control. Adjusted P values are <0.0001. (H) Bloating as measured by abdominal area of the fly in control, oenocyte specific Hnf4-i, Yki flies and Yki flies with oenocyte specific Hnf4-i. Adjusted P values are <0.0001. (I) *mElo* expression in oenocytes, normalized to housekeeping gene, in control, oenocyte specific Hnf4-I, Yki flies and in Yki flies with *Hnf4* knockdown. N = 2 biological replicates from 2 independent experiments. P values from left to right: <0.0001, 0.0002. (J) UMAP from whole body (without head) snRNAseq, *mElo* expression is enriched in adult oenocytes. (K) Quantification of pH3+ cell number, N =8 flies. P values are all <0.0001. (L-M) Lifespan analysis showing that oenocyte-specific knockdown of *Hnf4* (L) or *mElo* (M) in Yki flies, compared to only Yki or control flies, N = 40. Scale bar shows 20μm. P values are <0.0001. Data are presented as mean ± s.e.m. Statistical significance was determined using two-tailed Student’s t-test or one-way ANOVA with appropriate post hoc tests, or Log-rank (Mantel-Cox) test for lifespan assay. Ns is P > 0.05, *P < 0.05, **P < 0.01, ****P < 0.0001.

As a nuclear receptor, the localization of Hnf4 is highly enriched in oenocyte nuclei compared to the fat body [39, 46]. To visualize the localization of Hnf4, we tagged the endogenous *Hnf4* gene with 127D01, a small epitope tag that can be detected with a specific nanobody [47, 48], using an homology directed repair pathway CRISPR-Cas9 targeted insertion method [47]. To validate the Hnf4-127D01 line, we knocked down *Hnf4* in oenocytes, which significantly reduced its nuclear localization (Figure S1A). Hnf4-127D01 in control oenocytes showed high level of nuclear localization, consistent with previous findings (Figure 1E) [46]. Interestingly, increased nuclear localization of Hnf4-127D01 was observed in the oenocytes of Yki flies (Figure 1E-1F). Altogether, these data suggest that oenocytes in Yki flies have increased Hnf4 activity. To test whether oenocyte Hnf4 signaling is required for tumor growth, we measured ISC-mitosis using phospho-histone 3 (pH3+) staining. Downregulation of *Hnf4* in oenocytes of Yki flies *(Esg-LexA>yki^3SA^*-GFP*; PromE>Hnf4-i)* for 3 days showed a decrease in pH3+ staining compared to controls (*Esg> UAS-yki^3SA^*-*GFP*), and a slower growth rate for tumor (Figure 1G). Importantly, *Hnf4* knockdown in wild-type animals did not alter pH3+ cell numbers relative to controls (Figure 1G), indicating that the requirement for Hnf4 in Yki-driven tumors reflects its tumor-associated mis-regulation rather than a general role in proliferation. To quantify tumor burden, we measured the percentage of GFP-positive area in the gut and found that oenocyte-specific depletion of *Hnf4* in Yki flies markedly reduced GFP-positive area, consistent with tumor regression (Figure S1B-C). We next assessed tumor-induced cachexia phenotypes, including ovary wasting and abdominal bloating caused by fluid accumulation (Figure S1D). Short-term oenocyte *Hnf4* knockdown (3 days) alleviated ovary wasting (Figure S1E), reduced bloating (Figure 1H) and decreased whole-body wet weight (Figure S1F), supporting a role for aberrant oenocyte Hnf4 activity in promoting tumor growth and associated cachexia.

Our results suggest that Hnf4 transcriptional activity is induced in oenocytes of Yki flies. To validate this, we measured whether the expression of Hnf4 targets in Yki oenocytes also increased. Hnf4 target genes are involved in beta-oxidation and very long chain fatty acid (VLCFA) biosynthesis in oenocytes [39, 49]. Thus, we tested whether VLCFA biosynthesis genes (*Hnf4, mElo, CG30008, CG16904, FarO, ACC, HDAC, TER, Cyp4g1, Cpr*), as well as genes involved in beta-oxidation (*Whd, Scu, Yip2* and *CG7461)* were dysregulated in Yki flies. *CG30008, CG16904, FarO, ACC, HDAC, TER, Cyp4g1, Cpr, Whd, Scu, Yip2* showed no difference between control and Yki flies (Figure S1G). However, we observed a significant increase in *mElo* (Figure 1I), which encodes a methyl-branched cuticular hydrocarbon elongase [37], and a minor increase for *Hnf4 and CG7461* (Figure S1G).

mElo has recently been shown to produce long chain methyl-branched cuticular hydrocarbons in oenocytes that help prevent desiccation [37], a key function of Hnf4 in oenocytes [39]. Additionally, *mElo* expression is highly enriched in control adult oenocytes according to snRNAseq data (Figure 1J). Inhibition of *Hnf4* reduced *mElo* expression, and overexpression of *Hnf4* led to increased levels of *mElo* (Figure S1H-I), consistent with a previous report that *mElo* is a transcriptional target of Hnf4 [39]. Further*, mElo* is highly induced in the oenocytes of Yki flies and this induction is blocked when *Hnf4* is knocked down (Figure 1I).

In control flies, reduction of *mElo* in oenocytes did not affect pH3+ number in the gut (Figure 1K). To test the function of oenocyte *mElo* during tumor growth, we reduced the level of oenocyte *mElo* in Yki flies and observed fewer pH3+ in the gut, suggesting that *mElo* contributes to Yki tumor growth (Figure 1K). Similarly, we observed a reduced tumor burden with *mElo* knockdown in Yki flies (Figure S1J). We validated the effectiveness of *mElo* knockdown in the oenocytes (Figure S1K). Unlike *Hnf4,* reducing *mElo* expression in control oenocytes using the gene-switch driver (hereafter referred to as PromE^gs^) [30], where the driver is activated upon mifepristone feeding (RU), did not cause steatosis and affect ovary size, suggesting that other targets of Hnf4 regulate oenocytes steatosis and ovary size (Figure S1L and Figure S1M). Consistently, *mElo* knockdown did not rescue ovary wasting in Yki flies (Figure S1N).

To assess whether oenocyte lipid metabolism affects the viability of Yki flies, we inhibited *Hnf4* in oenocytes and observed a significant lifespan extension (Figure 1L).Inhibition of *mElo* in oenocytes of Yki flies also significantly increased lifespan (Figure 1M). Finally, mElo has a sex specific function in regulating different types of hydrocarbons [37]. Therefore, we tested whether mElo also regulates Yki viability in male flies. Strikingly, inhibition of *mElo* also significantly increased the lifespan in Yki males (Figure S1O).

### Oenocyte Hnf4 activity is regulated by TORC1

We next sought to determine the factors regulating Hnf4 level in oenocytes. Previous studies have shown that inhibiting the TORC1 pathway in oenocytes leads to steatosis [50], a phenotype that mimics reduced *Hnf4* expression in these cells [39], suggesting that the TORC1 pathway positively regulates *Hnf4* transcription in oenocytes. The Tuberous Sclerosis Complex (TSC) is known to negatively regulate TORC1 by inhibiting RAS homolog enriched in brain (Rheb) activity [51]. Next, we also tested the effects of oenocyte *TSC1,2* in tumor growth. Consistent with *Hnf4* and *mElo* knockdown, *TSC1,2* overexpression in oenocytes reduced tumor induced WE levels (Figure 2A). We also observed an increased level of TORC1 activity, as measured by phospho-4EBP (p4EBP) staining in oenocytes of Yki flies (Figure 2B-C). Additionally, overexpressing *TSC1,2* blocked *mElo* expression in oenocytes of Yki flies (Figure 2D), Yki-induced pH3^+^ cells in the gut (Figure 2E), as well as Yki-induced ovary wasting and bloating (Figure S2A and S2B). Together, these data identify oenocyte TORC1 activation as a key driver of tumor growth and implicate Hnf4 as a downstream effector.

**Figure 2:**
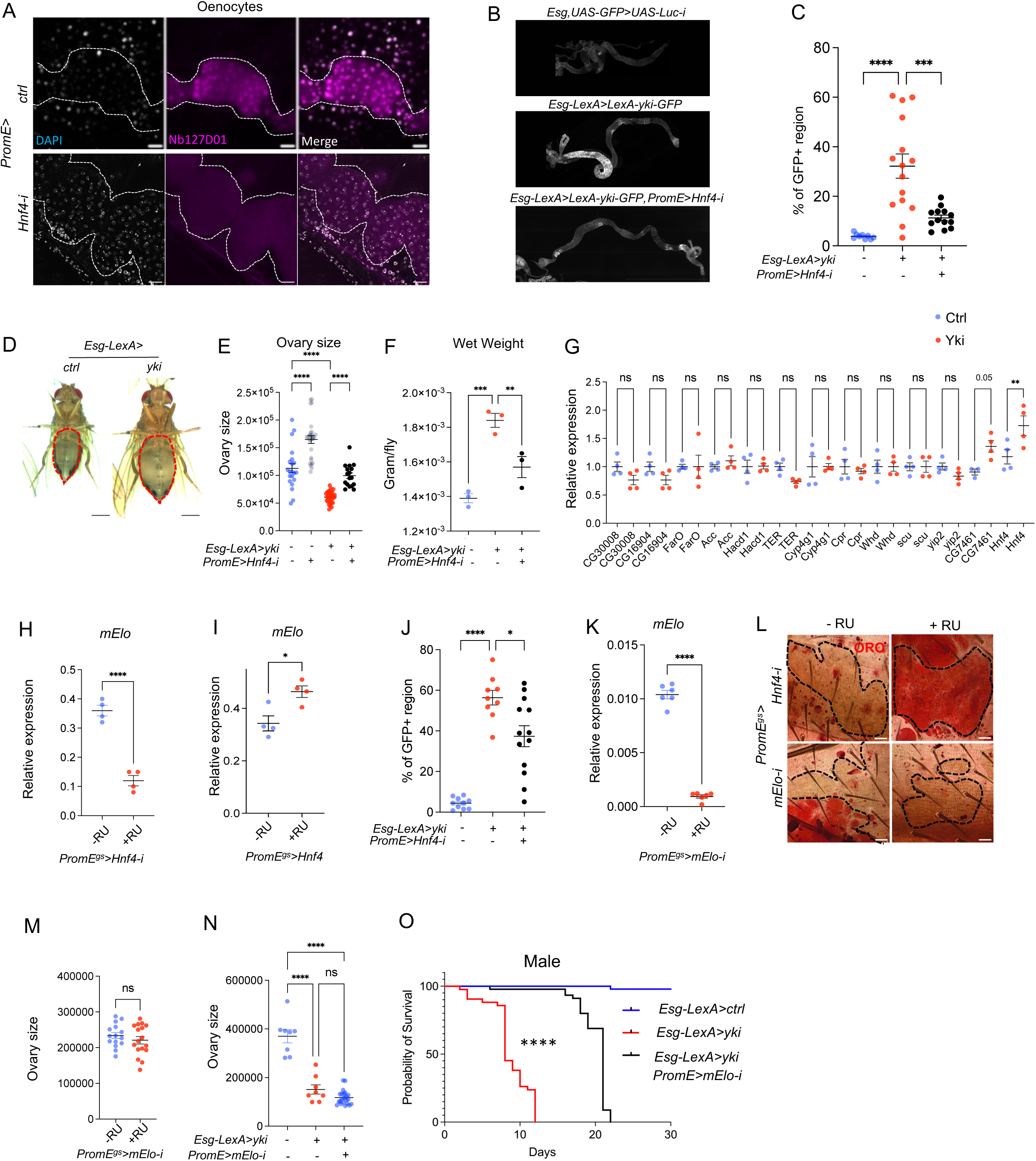
TORC1 regulates Hnf4 activity and lipid metabolism in oenocytes. (A) Whole body WEs level in ISC Yki flies and Yki with oenocyte *TSC1,2* overexpression. N = 3 biological replicates. P values from left to right: 0.0001, 0.0002. (B) Image of P-4EBP intensity is shown on the right (C), N = 18 flies in *Ctrl* and N = 15 flies in *yki*. P value is 0.0023. (D) *mElo* expression in whole body mRNA extracts in Yki flies and Yki flies with *TSC1,2* overexpression in oenocytes, N = 6 biological samples. P values are <0.0001. (E) Quantification of pH3+ cell number, sample sizes (left to right) were n = 9, 9, 13, 10. Adjusted P value <0.0001. (F) Violin plot of *Hnf4* expression in fat body, muscle and oenocytes upon overexpression of *TSC1,2* in oenocytes [12]. P value is <0.0001. (G-H) Immunostaining of Hnf4-127D01 in oenocytes when *TSC1,2* is overexpressed in oenocytes compared to control. DAPI marks nuclei, with dashed lines enclosing oenocytes, quantification of Hnf4-127D01 signal intensity in the nucleus, N = 8 flies. P value is <0.0001. (I) ModuleScore from snRNAseq analysis visualizes VLCFA synthesis gene expression across muscle, fat body and oenocytes, in control and upon oenocyte *TSC1,2* overexpression. (J) ORO staining indicates steatosis in oenocytes induced by *TSC1,2* overexpression, and in *TSC1,2* with *Hnf4* overexpression, N = 8. (K-L) Expression of Hnf4 targets *mElo* and *FASN2* in *TSC1,2* overexpression and in *TSC1,2* with *Hnf4* overexpression. N = 2 biological replicates from 2 independent experiments. P value is 0.0128 (K) and 0.0312 (L). Data represent mean ± s.e.m. Statistical significance was determined using two-tailed Student’s t-test or one-way ANOVA with appropriate post hoc tests, Ns is P > 0.05, *P < 0.05, **P < 0.01, ****P < 0.0001.

To test whether TORC1 regulates *Hnf4* expression level, we analyzed a previously published snRNAseq dataset [50]. Overexpression of *Tsc1* together with *Tsc2* (*TSC1*,2) in oenocytes, which inhibits *TORC1*, resulted in reduced *Hnf4* expression specifically in oenocytes (Figure 2F), but not in muscle nor fat body tissues, suggesting that TORC1 positively regulates Hnf4 levels in oenocytes. Next, we examined the protein level of endogenously tagged Hnf4-127D01 following *TSC1,2* over-expression and observed a significant reduction in Hnf4-127D01 levels in oenocytes, particularly in the nucleus as Hnf4 is a transcription factor (Figure 2G-H). Given that Hnf4 regulates hydrocarbon synthesis in oenocytes and its knockdown increases the susceptibility to dry starvation [39], we overexpressed *TSC1,2* in oenocytes and found that it led to an increased sensitivity of flies to dry starvation (Figure S2C). Additionally, as Hnf4 regulates the expression of genes involved in very-long-chain fatty acid (VLCFA) synthesis [39], we analyzed the expression of VLCFA synthesis genes in snRNAseq data from control (*PromE>EGFP*) versus TSC1,2-overexpressing oenocytes (*PromE>TSC1,2*). Using uniform Manifold Approximation and Projection (UMAP), which identifies cell types in mixed populations based on snRNAseq data, we found that VLCFA synthesis genes were highly enriched in oenocytes (Figure 2I). The ModuleScore, calculated by averaging the expression of the VLCFA synthesis pathway genes and subtracting the expression of a randomly selected gene set expression, decreased in oenocytes upon *TSC1,2* overexpression (Figure 2I).

Inhibition of TORC1 through overexpressing *TSC1,2,* or overexpressing Proline-rich Akt substrate 40 kDa (*PRAS40)*, or by feeding flies with rapamycin, all induced steatosis in oenocytes (Figure 2J and S2D-E), as observed following *Hnf4* knockdown in oenocytes (Figure S1L). However, simultaneous overexpression of *Hnf4* with *TSC1*,2 prevented the development of steatosis (Figure 2J), indicating that TORC1 regulates oenocyte lipid metabolism via Hnf4. Lastly, *TSC1,2* overexpression also reduced the expression of *mElo* and *FASN2,* which are Hnf4 target genes, and overexpressing *Hnf4* simultaneously blocked this decrease (Figure 2K-L). Together, these findings suggest that Hnf4 acts downstream of TORC1 to regulate lipid metabolism in oenocytes.

### ISCs secreted Pvf1 regulates Hnf4 through PvR signaling in oenocytes and aAects tumor associated phenotypes

TORC1 activity is regulated through various pathways, one of which includes PDGF/VEGF signaling. Studies have shown that PvR activation can stimulate the TOR pathway [50, 52, 53]. Consistent with this, muscle-secreted Pvf1 can regulate TORC1 activity in oenocytes, suggesting that the activation of the TORC1-Hnf4 axis in oenocytes of Yki flies may occur through PvR activation. Yki-expressing ISCs can secrete higher levels of the Pvf1 ligand, which activates PvR signaling in surrounding tissues (Figure S3A) [22].

However, *Pvf1* is expressed at low levels in oenocytes and is not significantly induced in oenocytes and muscles of Yki flies (Figure S3A-B). To determine whether tumor secreted Pvf1 non-autonomously activates Hnf4 level in oenocytes, we knocked down *Pvf1* expression in the ISCs of Yki flies (*Esg>yki^3SA^; Pvf1-i*) and observed its effect on Hnf4 nuclear level in oenocytes. *Esg>yki^3SA^; Pvf1-i* significantly reduced the level of Hnf4 nuclear localization compared to *Yki* flies (*Esg>yki^3SA^; Luc-i)* (Figure 3A-B). Consistently, *mElo* expression was reduced in *Esg>yki^3SA^; Pvf1-i* flies compared to Yki flies (Figure 3C). We also tested whether *Pvf1* reduction in Yki flies affects tumor development, like *Hnf4* inhibition in oenocytes. As reported previously [22], inhibition of *Pvf1* in the tumor gut rescues both bloating and ovary wasting (Figure 3D and Figure S3C). Gut tumors in *Esg>yki^3SA^;Pvf1-i* flies exhibited slower growth at early stage, as shown by reduced number of pH3+ cells and less GFP positive area (Figure 3E and Figure S3D).

**Figure 3:**
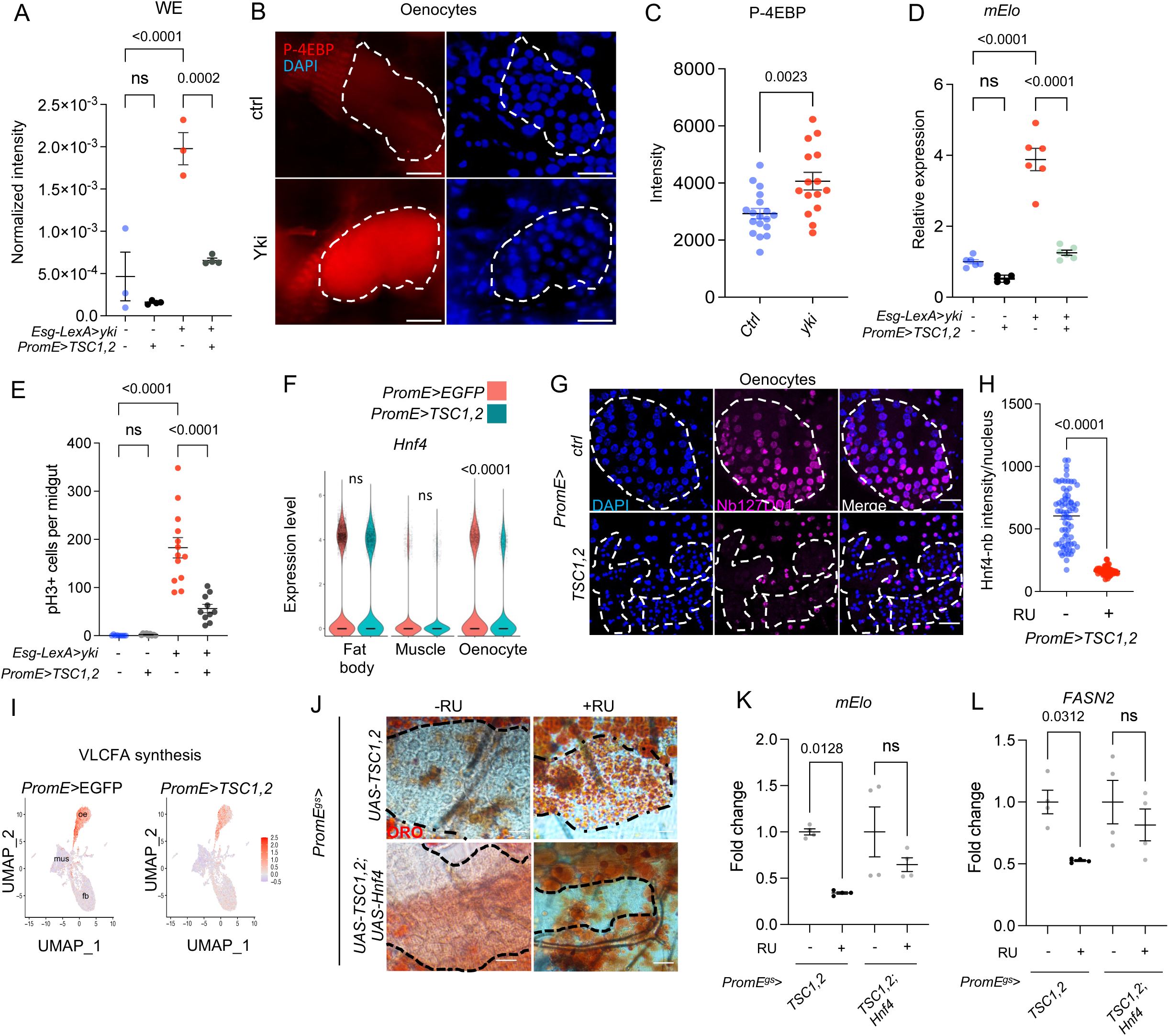
ISCs secreted Pvf1 activates Hnf4 through PvR signaling in oenocytes and aTects tumor associated phenotypes. (A-B) Hnf4 immunostaining in oenocytes of Yki flies with ISC-specific Pvf1 knockdown. Dashed lines enclose oenocytes. N = 10 biological replicates. P values are <0.0001. (C) *mElo* expression in oenocytes of Yki flies with ISC-specific Pvf1 knockdown, N = 4 biological replicates from 2 independent experiments. P values are 0.02034, 0.0074. (D) Inhibition of *Pvf1* in the gut affects bloating in Yki flies, sample sizes (left to right) were n = 19, 20, 22, 18 flies. P values from left to right: 0.001,0.0017. (E) ISC proliferation in Yki flies with *Pvf1* knockdown, sample sizes (left to right) were N = 21, 9, 10, 14 flies. Adjusted P values from left to right: <0.0001, 0.0079. (F) *mElo* expression in oenocytes of Yki flies with *PvR* knockdown. N = 2 biological replicates from 2 independent experiments. Adjusted P values from left to right: 0.0008, 0.0012. (G-H) *PvR* knockdown in oenocytes of Yki flies and its effect on tumor-associated bloating. N = 10 biological replicates. P value from left to right: <0.0001, 0.0071 (G) and pH3+ cell number, N = 10 biological replicates, P values from left to right: <0.0001, 0.0017 (H). (I) Lifespan extension in Yki flies with oenocyte-specific *PvR* knockdown, N = 40 flies. N = 10 biological replicates. P value <0.0001. (J-K) Hnf4 immunostaining in oenocytes of flies with ISC-specific *Pvf1* overexpression. Dashed lines enclose oenocytes. N = 10 biological replicates. P value < 0.0001 (K). (L) *mElo* expression in oenocytes following ISC-specific *Pvf1* overexpression. N = 4 biological replicates from 2 independent experiments. P value is 0.009. (M-O) Immunostaining and qPCR analysis reveal Hnf4 nuclear localization (N = 8 biological replicats) and *mElo* expression in oenocytes overexpressing active *PvR* (N = 3 biological replicates). P < 0.0001 (N) and P < 0.0001 (O). Dashed lines enclose oenocytes in all immunostaining images. Data are presented as mean ± s.e.m.; statistical significance was assessed using two-tailed Student’s t-test or ANOVA. Ns is P > 0.05, *P < 0.05, **P < 0.01, ****P < 0.0001.

To demonstrate that Pvf1 secreted from Yki tumors directly controls *mElo* expression through PvR, we used the dual Gal4 and LexA systems to reduce *PvR* level in oenocytes while inducing Yki in ISCs. *Esg-LexA>yki^3SA^;PromE>PvR-i* animals showed reduced *mElo* expression (Figure 3F), decreased the level of bloating (Figure 3G) and ovary wasting (Figure S3E), reduced the number of pH3+ cells in gut tumors (Figure 3H) and reduced tumor burden (Figure S3F), and extended lifespan compared to Yki flies (Figure 3I). Altogether, these data suggest that PDGF/VEGF induced by Yki tumor non-autonomously increases Hnf4 activity in oenocytes through PvR.

To test whether ISC secreted Pvf1 can induce oenocyte Hnf4 levels under normal physiology, we overexpressed *Pvf1* specifically in adult ISCs (*Esg>Pvf1*) and measured Hnf4 nuclear level in oenocytes. *Pvf1* overexpression in ISCs increased nuclear Hnf4 levels (Figure 3J-K), suggesting an increased transcriptional activity as measured by increased level of *mElo* expression (Figure 3L). To test whether ISC-derived Pvf1 activates PvR in oenocytes, we overexpressed an active form of *PvR* in oenocytes which was associated with increased nuclear Hnf4 level, as well as increased level of *mElo* expression (Figure 3O).

### Circulating lipids target trachea but not ISCs to promote trachea growth

To identify the lipid species induced by oenocytes in Yki flies, we performed lipidomic analysis on the fly hemolymph of control flies, *Esg-LexA> yki^3SA^* and *Esg-LexA> yki^3SA^; PromE>Hnf4-i* flies (Figure 4A, Supplementary Data 3). Among the lipid species that are induced by Yki and decreased by *Esg-LexA > yki^3SA^; PromE>Hnf4-i*, the majority are phospholipids including phosphotidyl choline (PC), phosphatidylglycerol (PG) and phosphatidylethanolamine (PE), triglycerides (TGs) and acylcarnitines (AcCa). More specifically, AcCa 24:1 and AcCa 24:2, which are very-long-chain acylcarnitines (C≥22) secreted by mitochondria when metabolism is inhibited, were elevated in Yki flies while reduced by *Hnf4* knockdown (Figure 4B-C). Circulating PC 36:4, and PE 32:5 phospholipids and TG(18:3_16:0_16:1), TG(18:2_16:0_18:0) triglycerides were induced in Yki flies and rescued by *Hnf4* knockdown (Figure 4D-G). Interestingly, PC 36:4 is elevated in the serum and plasma samples from breast cancer patients and TG(18:2_16:0_18:0) is induced in bladder cancer patients [54].

**Figure 4:**
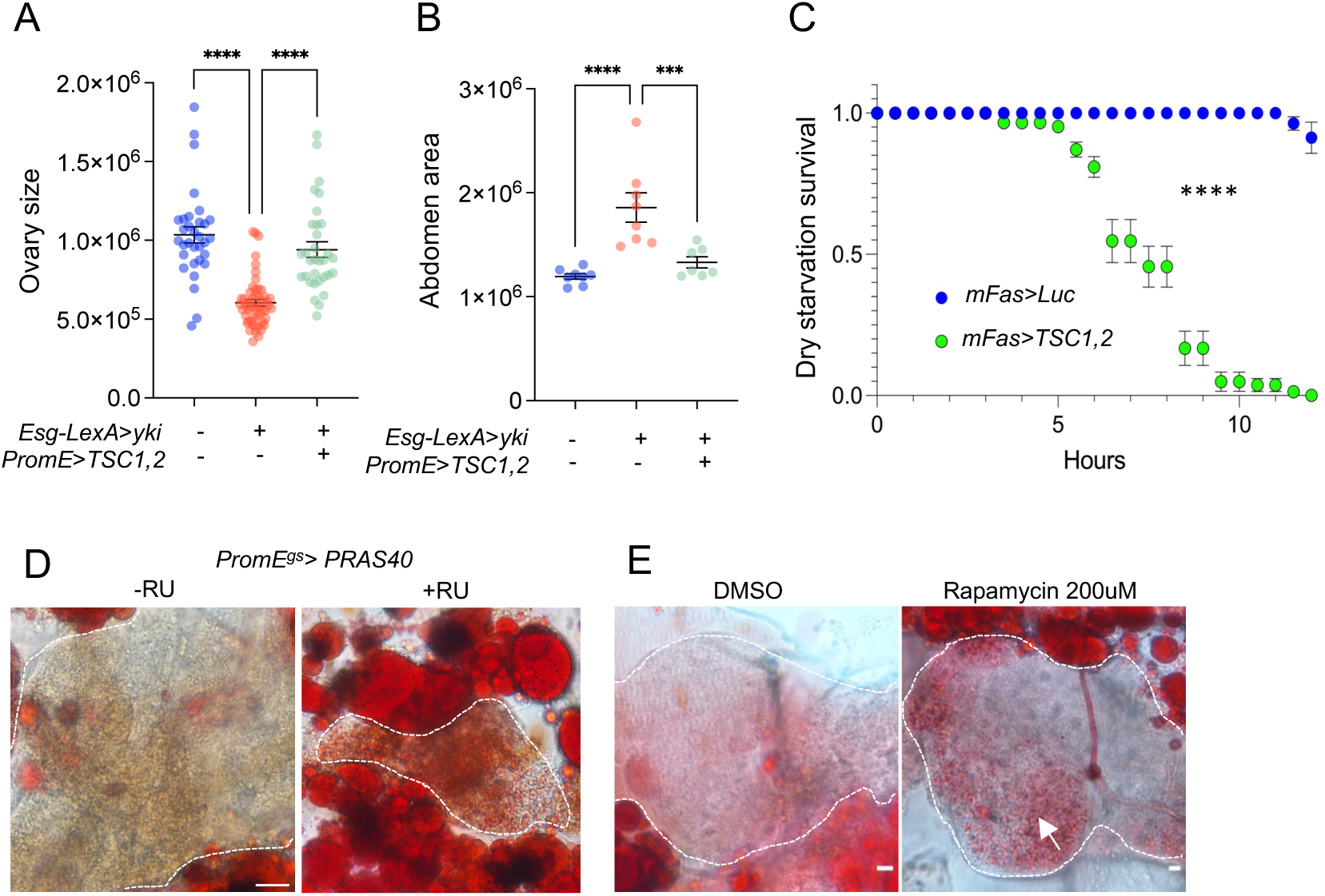
Circulating lipids target trachea but not ISCs to promote tracheal growth. (A) Untargeted lipidomic analysis of fly hemolymph showing lipid species in control, Yki flies, and Yki flies with oenocyte-specific *Hnf4* knockdown (*Esg>yki^3SA^;PromE>Hnf4-i*). N = 4 biological replicates. For the full names corresponding to the abbreviations, see Supplementary Data 4. (B-C) Levels of very-long-acylcarnitines (AcCa 24:1 and AcCa 24:2) in Yki flies with or without *Hnf4* knockdown in oenocytes. Adjusted P value from left to right: 0.0185, 0.0415 (B). Adjusted P value = 0.0054 (C). (D-E) Levels of circulating phospholipids, including PC 36:4 PE 32:5, across experimental conditions. N = 4 biological replicates. Adjusted P value is 0.0018, 0.0381 (D). P values <0.0001 (E). (F-G) Levels of circulating TG 18:3_16:0_16:1 and TG 18:2_16:0_18:0 in the fly hemolymph. N = 4 biological replicates. Adjusted P value from left to right is 0.0281, 0.0281 (F). P value is 0.026 (G). (H-J) Number of tracheal branches, tube area, and skeleton length in Yki flies with ISC-specific knockdown of *LpR1* and *LpR2*. Sample sizes (left to right) were N = 14, 16, 11, 16. Adjusted P value is 0.0001 (H). Adjusted P is 0.0021 (I). Adjusted P <0.0001 (J). (K) PH3+ cell counts in ISCs of Yki flies with ISC-specific knockdown of *LpR1* and *LpR2*. N = 8-10 flies. P values from left to right: 0.0257, 0.0001, 0.013. (L) PH3+ cell counts in ISCs of Yki flies with trachea-specific (*Btl-Gal4* driver) knockdown of *LpR1* and *LpR2*. P values from left to right: <0.0001, 0.0118, 0.0323. (M) GFP positive region in the gut of tracheal specific *LpR2* knockdown in Yki flies. N from left to right is 8, 8, 9, 5. P value from left to right: <0.0001, 0.0004. (N) Abdominal bloating area in Yki flies with trachea-specific LpR2 knockdown. P value from left to right: <0.0001, <0.0001, 0.0487. (O-Q) Number of tracheal branches, tube area, and skeleton length in Yki flies with trachea-specific knockdown of *LpR2*. N = 8-15 biological replicates. Adjusted P from left to right: <0.0001, <0.0001, 0.0094 (O). Adjusted P from left to right: <0.0001, <0.0001, 0.0048 (P). Adjusted P from left to right: <0.0001, <0.0001, 0.011 (Q). Data are presented as mean ± s.e.m.; statistical analysis was performed using two-tailed Student’s t-test or ANOVA as appropriate. Ns is P > 0.05, *P < 0.05, **P < 0.01, ****P < 0.0001.

PC and PE are major membrane phospholipids and contribute to lipoprotein assembly and lipid transport. Thus, elevated PC/PE could promote tracheogenesis by supporting both membrane biogenesis and lipid delivery needed for tracheal membrane expansion. Increased AcCas and WEs could contribute to tracheogenesis by supporting local energetic demand or increasing the production of reactive oxygen species (ROS). For example, hepatocyte-derived circulating AcCas are required for systemic thermogenesis during cold exposure in mice; in this setting, peripheral tissues—including brown adipose tissue, skeletal muscle, and heart—take up plasma acylcarnitines and oxidize them to support energy metabolism [55]. In insects, lipid transport in circulation primarily relies on lipophorins (Lpps), which are the major lipoproteins. It is produced predominantly by the fat body and delivers neutral lipids to peripheral tissues via the lipophorin receptors LpR1/LpR2, which mediate lipoprotein binding and internalization through endocytic pathways [56, 57]. The increased circulation of PC 36:4 and PE 32:5 in Yki flies suggests an increase in lipoproteins that transport lipids from oenocytes towards trachea or the midgut.

The transport of lipids, such as phospholipids, from oenocytes to the trachea or ISCs to promote tumor growth has not been reported [42]. To test whether changes in oenocyte metabolism regulates ISCs growth, we inhibited Hnf4 activity in oenocytes by overexpressing *TSC1,2*, knocking down or overexpressing *Hnf4*, and observed no significant effect on the number of pH3+ cells in the midgut (Figure S4A). This suggests that changes in oenocyte lipid metabolism does not promote ISC proliferation under normal condition.

Since 95% of the hemolymph lipids are transported through lipophorin (Lpp) in flies [56], and that low-density lipoprotein (LDL) receptor homologues LpR1 and LpR2 promote Lpp uptake [58], we reduced *LpR1* or *LpR2* expression level in the ISCs. However, this had no impact on the trachea area, skeleton length, and only a minor increase in the number of branches in the *LpR1* knockdown condition (Figure S4C-D). Reducing LpR1 and LpR2 activities in the ISCs also did not affect the proliferation (Figure S4E). To validate RNAi knockdown efficiency, we quantified *LpR1* expression in thoraces from flies expressing *LpR1* RNAi under the muscle-specific *Mhc-Gal4* driver (Myosin heavy chain), and *LpR2* expression in fat bodies from flies expressing *LpR2* RNAi under the fat body–specific *Lpp-Gal4* driver (Apolipophorin). Both RNAi lines efficiently reduced their respective target transcripts (Figure S4F-G ). To further examine the role of LpR1 or LpR2 in ISC tumors, we reduced *LpR1* and *LpR2* levels while overexpressing Yki (*Esg> yki^3SA^;LpR1-i and Esg> yki^3SA^;LpR2-i*), and found that these manipulations did not impair Yki-induced tracheal growth (Figure 4H-J) or prevent Yki-induced ISC proliferation (Figure 4K). These results suggest that ISC proliferation does not rely on circulating lipids in the hemolymph and that circulating lipids control tracheaogenesis instead.

We further tested whether disrupting *LpR1* and *LpR2* activities in the trachea could affect tumor growth. It has been reported that restricting tracheal growth delays tumor development [42]. Consistently, knocking down *LpR2*, but not *LpR1*, using a temperature sensitive trachea-specific driver (*btl-GAL4, UAS-srcGFP, Tub-GAL80^TS^,* referred to as *Btl*) in Yki flies prevented Yki-induced ISC proliferation (Figure 4L) and the tumor burden (Figure 4M). Consistently, it also reduced the level of bloating (Figure 4N), tracheal number of branches, tube area and skeleton length compared to control flies (*Btl>+*) (Figure 4O-Q). Knocking down *LpR2* in the normal condition also reduced tracheal branch number, skeleton length and total tube area (Figure 4O-Q). Reducing *LpR1* and *LpR2* function in the trachea did not alter ISC proliferation under basal conditions (Figure S4F). The different phenotypes observed between *LpR1* and *LpR2* knockdowns might reflect differences in RNAi efficacy, as the *LpR2* RNAi line shows stronger knockdown than the *LpR1* RNAi line (Fig. S4F–G). In addition, *LpR2* expression is modestly higher in tracheal cells of *Yki* flies at day 8, which could explain why *LpR2* depletion produces a more pronounced phenotype.

Interestingly, oenocyte-specific depletion of *LpR2* also altered gut tracheal morphology under basal conditions (Figure S4H-J), reflecting that the trafficking of phospholipids, WEs and AcCas from oenocytes relies on the uptake of LPPs. Altogether, these findings are consistent with a model in which tracheal morphology, rather than ISC proliferation, depends on lipids generated or processed in oenocytes and delivered via circulating LPP particles.

### Hnf4-mElo axis in oenocytes mediates lipid metabolism and tracheogenesis induced by Yki tumors

Tumor growth in the adult *Drosophila* midgut has been previously shown to promote tracheole density [42]. Similarly, we observed an increase in the number of trachea branches, total tube area, and total tube length in the surrounding tissue of the Yki midgut (Figure S5A-D). This led us to investigate whether oenocyte lipid metabolism mediated by Hnf4-mElo axis is crucial for trachea growth in the Yki midgut.

To explore whether perturbations of lipid metabolism in oenocytes could block Yki-induced tracheogenesis in the midgut, we manipulated lipid metabolism by overexpressing *TSC1,2* or knocking down *Hnf4* or *mElo*. Using the LexA/LexAop and Gal4/UAS systems, overexpressing *TSC1,2* or reducing *mElo* significantly decreased the number of tracheal branches, total tube area, and total tube length in the Yki midgut. Reducing *Hnf4* levels, on the other hand, only decreased the total tube area (Figure 5A-C).

**Figure 5:**
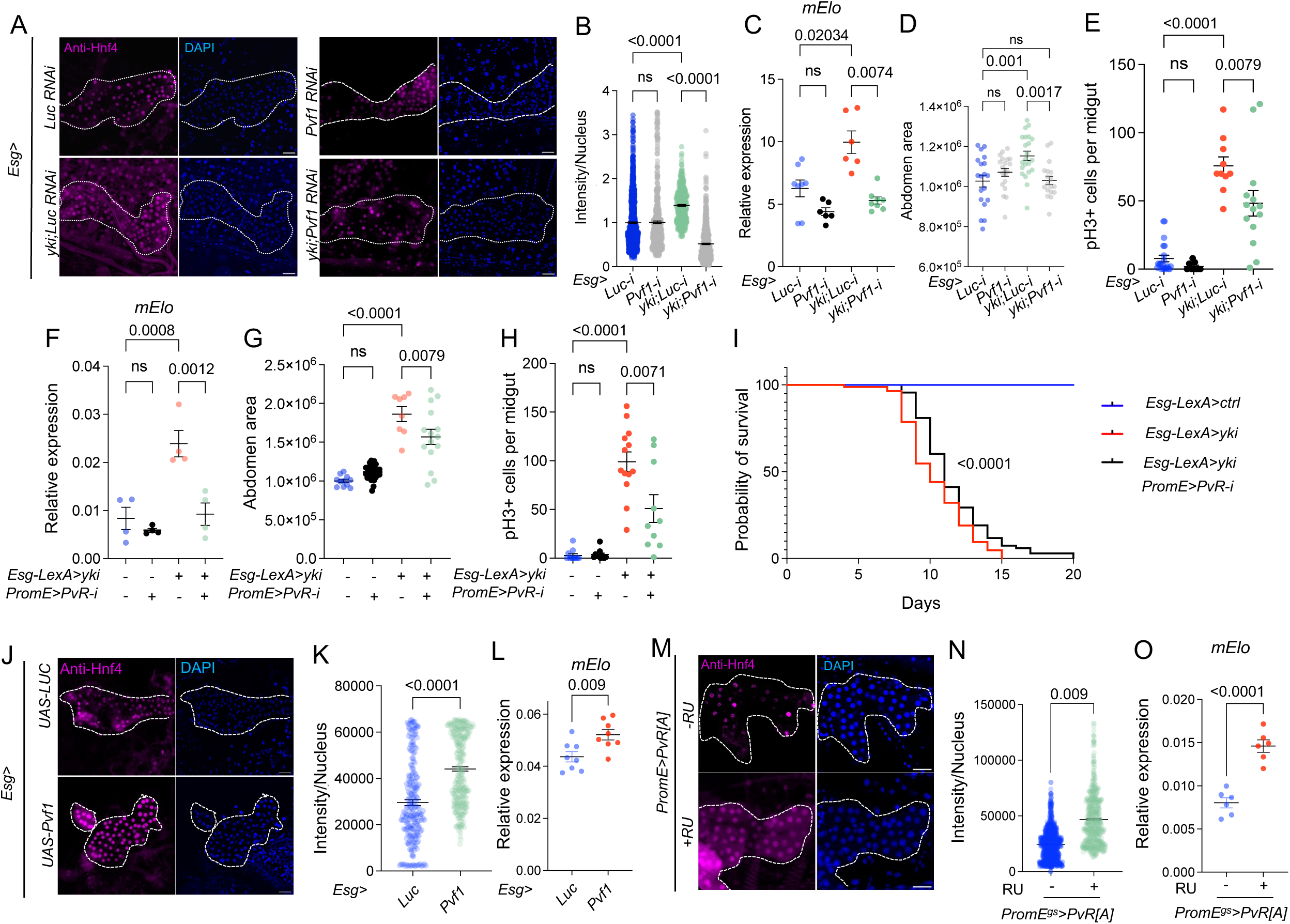
Hnf4-mElo axis in oenocytes mediates lipid metabolism and tracheogenesis induced by Yki tumors. (A-C) Number of tracheal branches, total tube area, and total tube length in the midgut of Yki flies with oenocyte-specific *TSC1,2* overexpression, *Hnf4* knockdown, or *mElo* knockdown. Sample sizes (left to right) were n = 15, 17, 10, 9, and 11 flies. Adjusted P values from left to right: 0.0303, 0.0002, 0.0004 (A). Adjusted P values from left to right: 0.0318, <0.0001, 0.0072, 0.0066 (B). Adjusted P values from left to right: 0.0276, 0.0005, ns, 0.0002 (C). (D-F) Tracheal morphology measurements, including number of branches and skeleton length, in flies with ISC-specific knockdown of *Pvf1* combined with *yki* overexpression (*Esg>yki^3SA^;Pvf1-i*). Sample sizes (left to right) were n = 11, 11, 9 flies. Adjusted P values from left to right: 0.008, 0.0029 (D). Adjusted P values from left to right: 0.0041, 0.0009 (E). Adjusted P values from left to right: 0.0438, 0.0022 (F). Dashed lines enclose oenocytes in all relevant images. Data are presented as mean ± s.e.m.; statistical analysis was performed using two-tailed Student’s t-test or ANOVA as appropriate. Ns is P > 0.05, *P < 0.05, **P < 0.01, ****P < 0.0001.

In addition, to verify whether midgut secreted Pvf1 regulates tracheal growth, we knocked down *Pvf1* in ISC in addition to Yki overexpression. Reduction of *Pvf1* blocked Yki induced number of branches, tube area and skeleton length (Figure 5D-F), indicating that oenocyte lipid metabolism is important for tracheal growth induced by Yki.

### Oenocyte Hnf4 and mElo regulate tracheogenesis in the midgut

Next, we aimed to understand whether oenocyte lipid metabolism controls trachea growth in control guts. Consistent with *Hnf4* knockdown, oenocyte-specific *mElo* reduction and *TSC1,2* overexpression showed reduced level of whole body WE in non-Yki flies (Figure 6A-B). WEs are essential parts of the outer layer of the trachea [59–62], and *waterproof* (*wat*), which is involved in biosynthesis of wax, has been implicated in hydrophobic tracheal coating in *Drosophila* embryos [62]. Knocking down *Hnf4* did not affect the transepithelial barrier mediated by septate junctions in the midgut trachea on the midgut, as revealed by red dextran injection experiment (Figure 6C). However, reduction of oenocyte *Hnf4* altered the morphology of trachea associated with the midgut (R2-R4 regions) (Figure 6D). Specifically, we observed a reduction in the number of branches and total skeleton length, normalized to the area of the gut surface (Figure 6E-G). Consistently, we also observed a reduction in branch numbers, tube area and skeleton length when *mElo* was knocked down in oenocytes of control flies (Figure 6H-J). The effect of oenocyte Hnf4 on trachea number is not specific to the midgut, as we also observed a reduction of trachea total tube area in the ovary of control flies (Figure S6A-D). We also examined whether Hnf4 and mElo in the fat body contribute to tracheal growth, since the fat body is a major metabolic organ involved in lipid homeostasis and, like oenocytes, shares hepatocyte-like functions in flies [63, 64]. However, knockdown of *Hnf4* and *mElo* using the fat body– and gut-specific GeneSwitch driver *S106-Gal4* [65] did not affect tracheal growth (Figure S6E-J ), in contrast to the marked reduction observed upon oenocyte-specific knockdown. It suggests that oenocytes play a more prominent role than the fat body in regulating tracheal growth. Overexpressing *Hnf4* or *mElo* is sufficient to induce total tracheal tube area and skeleton length in the midguts of control flies (Figure 6K-P). Finally, to eliminate the effect of RU feeding on trachea morphology, we fed control (*PromE*>*ctrl*) flies and observed no difference in trachea morphology (Figure S6K-M).

**Figure 6:**
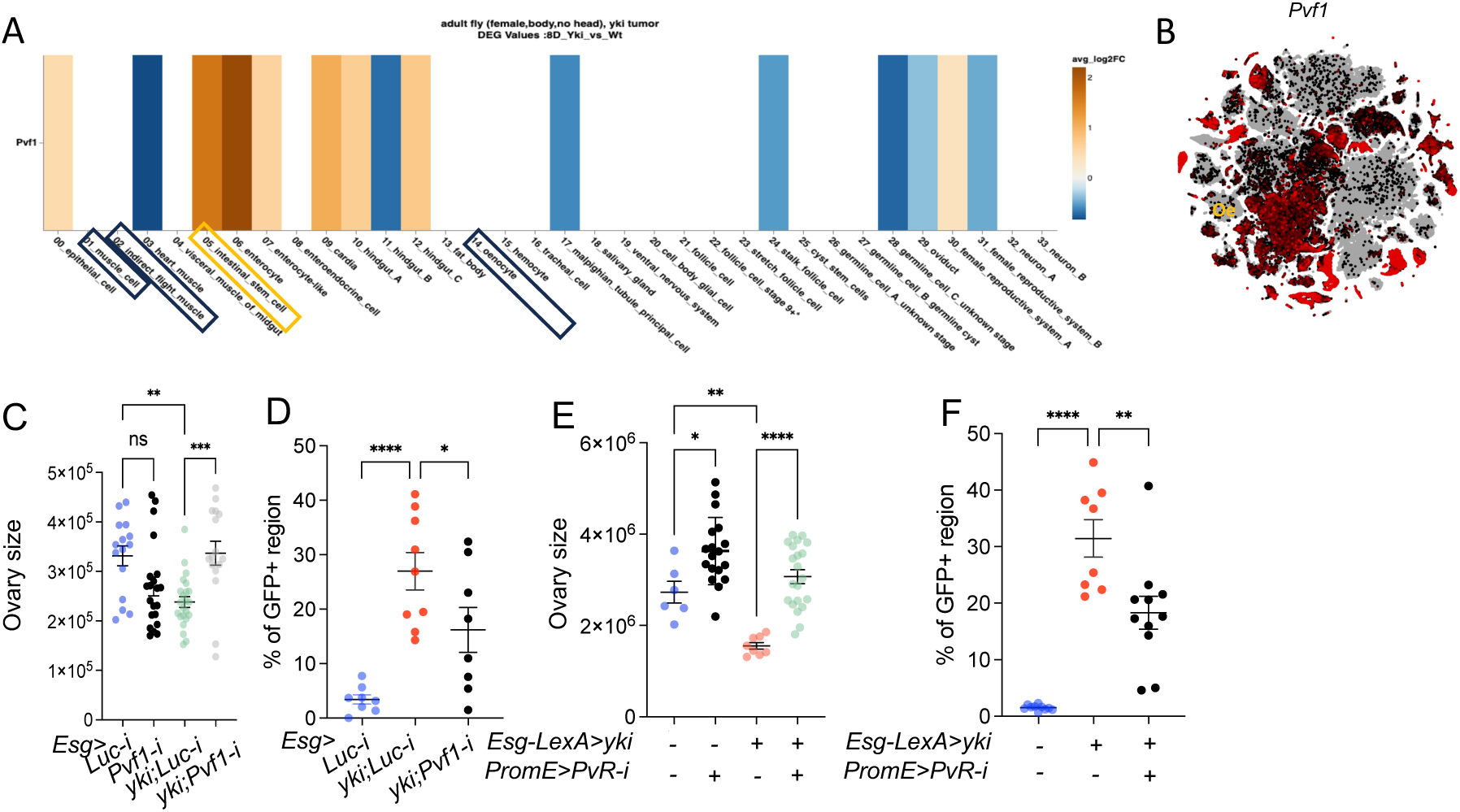
Hnf4-mElo axis regulates tracheogenesis in the midgut. (A-B) Whole-body wax ester (WE) levels in flies with oenocyte-specific *Hnf4* knockdown (*PromE^gs^>Hnf4-i*), *mElo* knockdown (*PromE^gs^>mElo-i*), or *TSC1,2* overexpression (*PromE^gs^>TSC1,2*). N = 3 biological replicates (A), N = 4 biological replicates (B). P value is 0.0177 (A). P value is 0.0005 (B). (C) Transepithelial barrier function of the trachea on the midgut, assessed using red dextran injection, in flies with oenocyte-specific *Hnf4* knockdown. N = 8 flies. (D-G) Trachea morphology on the midgut (R2-R4), including branch number and skeleton length normalized to gut surface area, in flies with *Hnf4* knockdown in oenocytes. N = 8 flies. P value is 0.0029 (E), ns (F), 0.005 (G). (H-J) Branch number, tube area, and skeleton length of the midgut trachea in flies with oenocyte-specific *mElo* knockdown. N = 6 flies. P value is 0.0395 (H), 0.0299 (I), 0.0203 (J). (K-P) Number of branches, tracheal tube area and skeleton length of the midgut in flies overexpressing *Hnf4* or *mElo* in oenocytes (*PromE^gs^>Hnf4* or *PromE^gs^>mElo*). Sample sizes (left to right) were n = 15, 8 flies. P value is 0.01869 (L), 0.0021 (M). P value is 0.0437 (O), 0.0323 (P). Data represent mean ± s.e.m, statistical significance was assessed using two-tailed Student’s t-test or ANOVA. Ns is P > 0.05, *P < 0.05, **P < 0.01, ****P < 0.0001.

### VEGF-A reprograms lipid metabolism in hepatoma cells, while hepatic lipid metabolism is altered in tumor-bearing mice

Using the *Drosophila* Yki tumor model, we found that the tumor secreted PDGF/VEGF-like factor Pvf1 can non-autonomously regulate Hnf4 activity in hepatocyte-like cells to promote the production of VLCFA or related lipids. To test whether this mechanism is conserved in mammals, we incubated HepG2 cells with vascular endothelial growth factor A (VEGF-A), which is orthologous to Pvf1 in flies and a potent pro-angiogenic growth factor expressed in many types of tumors. We measured the expression level of ELOVL fatty acid elongase 1 – 7 (*ELOVL1-7*). Incubating HepG2 with VEGF-A for 24 hours can significantly increase the level of *ELOVL1, 2, 4, 6* and *7*, suggesting a conserved regulation of PDGF/VEGF on lipid metabolism in hepatocytes (Figure 7A-E). HNF4*α* plays an important role in hepatocyte lipid metabolism. Consistent with this, *APOB*, an early transcriptional target of HNF4*α* [66, 67], showed increased expression in HepG2 cells following VEGF-A treatment (Figure 7F), suggesting an increased HNF4*α* activity.

**Figure 7:**
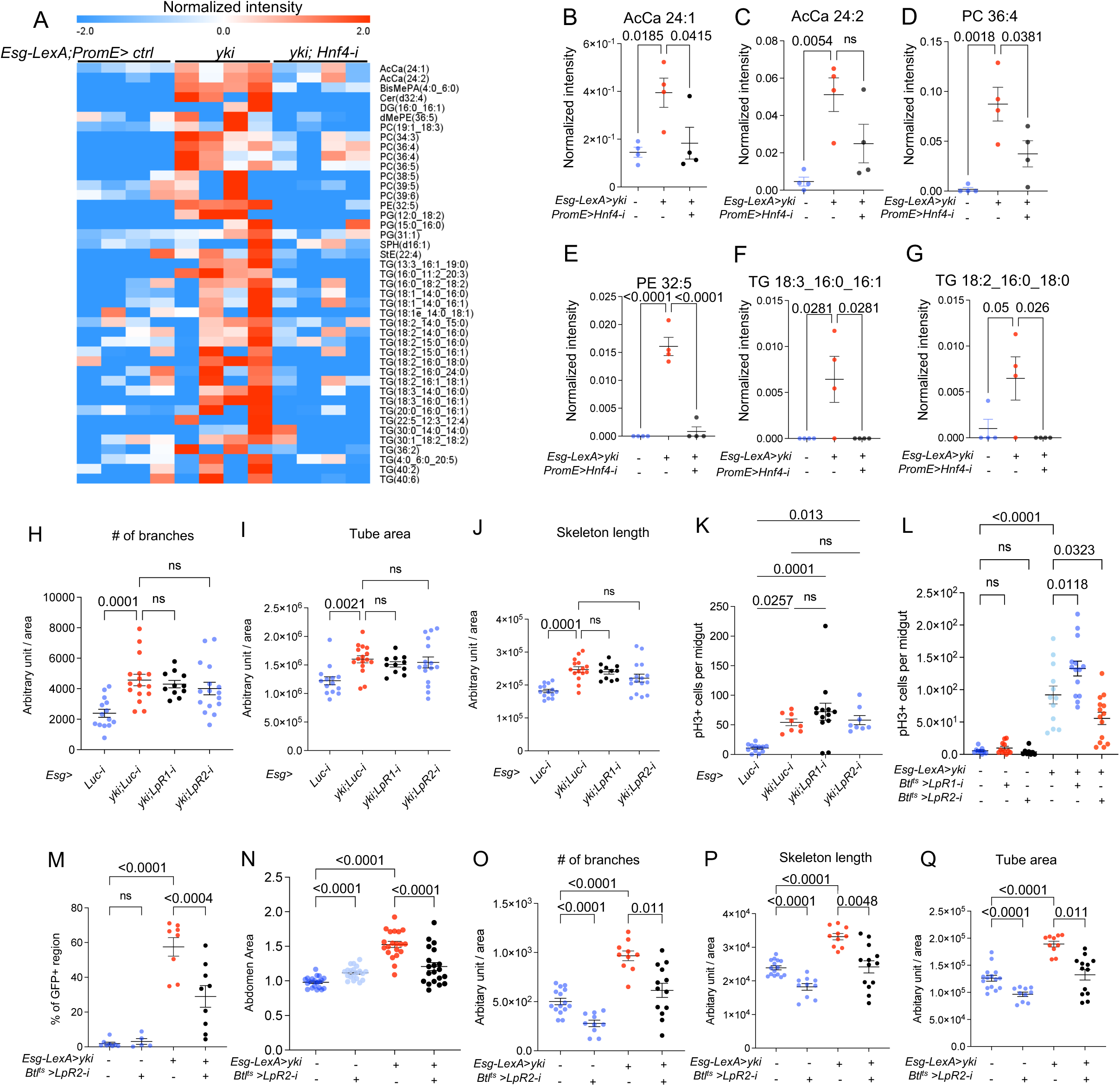
Regulation of hepatocyte lipid metabolism by pro-angiogenic factor in KL mice tumor model. (A-F) Relative expression levels of *ELOVL* fatty acid elongase 1,2,4, 6 and 7 and *APOB* in HepG2 cells incubated with vascular endothelial growth factor A (VEGF-A) for 24 hours, compared to controls. N = 4 biological replicates, from 2 independent experiments. P values are: 0.0096 (A), 0.0103 (B), 0.002 (C), 0.001 (D), 0.0032 (E), 0.0023 (F). (G) The level of WE from conditioned medium collected from HepG2 cultures following VEGF-A treatment. N from left to right is 3, 4 biological replicates. P value is 0.002. (H-L) Relative expression levels of *Hnf4α, Elovl7, Acc1* and *Fasn* in liver samples from *KrasG12D/+;Lkb1f/f* (KL) tumor-bearing mice compared to control mice. N = 3 biological replicates, from 2 independent experiments. P values are 0.0372 (H), 0.0377 (I), 0.0004 (J), 0.0044 (L). (M) The level of WEs from serum samples of KL mice and control mice. P value is 0.048 (M). (N) Heatmap showing the significantly changed lipid species from serum samples of tumor-bearing and control mice. For the full names corresponding to the abbreviations, see Supplementary Data 4. Fed is fed mice samples, cacs means cachexic tumor bearing mice samples. (O) Schematic representation of the proposed mechanism, depicting tumor-secreted PDGF/VEGF-like factors activating the mTOR-Hnf4 axis in hepatocyte-like oenocytes to promote lipid synthesis, which are then transported to trachea. Figure is created in BioRender. Huang, K. (2026) https://BioRender.com/9vaa42y. Data are presented as mean ± s.e.m.; statistical analysis was performed using two-tailed Student’s t-test and significance analysis of microarrays was used for lipid species. Ns is P > 0.05, *P < 0.05, **P < 0.01, ****P < 0.0001.

Consistently, we observed an increased level of HNF4*α* localization in HepG2 cells treated with VEGF-A (Figure S7A-B). We performed lipidomic profiling of conditioned medium collected from HepG2 cultures following VEGF-A treatment and observed a robust increase in WE (Figure 7G, Supplementary Data 7), as seen in Yki flies (Figure 1A-C).

Together, these results suggest a conserved VEGF-A–driven regulatory mechanism in hepatic lipid metabolism mediated by HNF4α.

In addition, we collected liver samples from *KrasG12D/+;Lkb1f/f* (KL) male mice, in which tumors were specifically induced in the lungs. The age of the mice ranged from 7-38 weeks. We analyzed the expression levels of *Hnf4α*, *Elovl1-7* and HNF4*α* target genes *acetyl-Coenzyme A carboxylase alpha* (*Acaca*), *fatty acid synthase* (*Fasn*) in the liver [68–70]. *Hnf4α, Elovl7, Acc1* and *Fasn* showed higher expression level in the tumor bearing mice, compared to control (*LKB1 f/f*), suggesting that tumor growth in the lung increased

HNA4*α* activity and induced genes involved in lipid metabolism in liver tissues in the KL mouse model (Figure 7H-L). Consistent with the expression data, lipidomic profiling of serum samples with tumor bearing male mice and control mice revealed higher level of WE in the tumor mice serum (Figure 7M), which are mostly comprised of very long chain fatty acids (Figure S7C). The lipidomic analysis also showed that out of 147 significantly changed lipid species, most of them (118) are induced and only 29 are reduced (Figure S7C). From the summed lipid species, we found that although phosphatidylcholine (PC) and triglycerides (TG) are reduced in tumor bearing mice serum, most other lipid species, such as Acylcarnitine (AcCa), lysophosphatidylcholine (LPC) are highly induced in tumor bearing mice (Figure 7N). Altogether, these data suggest a conserved mechanism of lipid metabolism regulation in hepatocytes by a PDGF/VEGF-like factor and Hnf4 (Figure 7O).

## Discussion

We reveal a mechanism by which gut tumors in *Drosophila* reprogram lipid metabolism in oenocytes to promote tracheal development and tumor growth. Our findings demonstrate that tumors secrete a PDGF/VEGF-like factor, Pvf1, which activates TORC1 activity, triggering the expression of *mElo* through Hnf4 in oenocytes. This TORC1-Hnf4 signaling pathway enhances the production of specific lipids, such VLCFAs and wax esters (WEs), which stimulate tracheal growth, a crucial process for tumor expansion. We also show that reducing Hnf4 or mElo activity in oenocytes suppresses tumor growth, tracheogenesis, and tumor-induced cachexia, while extending lifespan. Additionally, we highlight the conservation of this regulatory pathway in mammals, where VEGF-A stimulates lipid metabolism gene expression in human hepatocytes. Furthermore, lung tumor-bearing mice exhibit increased *Hnf4α* and *Elovl7* expression in peripheral hepatocytes.

Pvf1, a tumor-secreted factor, has previously been linked to tumor-induced muscle and fat body wasting, as well as renal dysfunction in *Drosophila* [22, 71, 72]. In this study, we uncover a role for tumor-secreted Pvf1 in modulating oenocyte lipid metabolism through TORC1 and Hnf4, which regulates the expression of the oenocyte-enriched gene *mElo* (Figure 3). While Hnf4 activity is known to be induced by starvation and during the developmental transition from pupae to larvae [39, 49], its upstream regulators remain poorly understood. In MEF cells, the HNF4*α* binding motif is highly enriched in the promoters of TORC1-induced genes [73], suggesting HNF4*α* mediates the transcriptional activities of TORC1. Here, we demonstrate that TORC1 positively regulates Hnf4 protein levels in the oenocyte nucleus through transcriptional regulation (Figure 2G). However, the precise mechanisms controlling *Hnf4* transcription downstream of TORC1 remain unclear. REPTOR and REPTOR-BP are likely key players in this process, as they are the major effectors of the transcriptional response induced by TORC1 inhibition [74].

Previous studies showed that muscle-derived Pvf1 suppresses lipid synthesis in *Drosophila* oenocytes and limits fat accumulation in the fat body through PI3K/Akt/TOR signaling [50]. In contrast, our results indicate that Yki tumor–derived Pvf1 promotes very-long-chain fatty acid synthesis in oenocytes, thereby supporting tumor growth. This difference may reflect tissue-specific effects of Pvf1. Notably, muscle-derived Pvf1 was reported to inhibit oenocyte lipid synthesis, whereas Pvf1 from the fat body and midgut had no detectable effect on oenocyte lipid metabolism or steatosis [50]. What a gut tumor–derived Pvf1 regulates oenocyte lipid metabolism has not been previously examined. One possible explanation is that tissue-specific post-translational modification of Pvf1 alters its activity, as PDGF/VEGF ligands function as disulfide-linked dimers [75, 76]. In addition, Pvf2 and Pvf3 are strongly induced in gut tumors [22], raising the possibility that distinct PVF homo- or heterodimers may generate different downstream signaling outputs [53].

Our study identified a mode of communication between oenocytes and the gut.

Oenocytes produce lipid substrates that support gut tracheal growth, a process crucial for tumor expansion and potentially ISC growth. Like tumors, bacterial infections and epithelial damage also trigger ISC proliferation [77–79], suggesting that oenocyte-secreted lipids may also support stress-induced ISC expansion. Lipids from oenocytes are possibly acquired by the tracheal cells instead of ISCs, as knockdowns of *LpR1* and *LpR2* in ISCs have no effects in tumor growth (Figure 4H-J). In contrast, reducing *LpR2* in tracheal cells was sufficient to decrease tracheal development (Figure 5O-Q). This may be due to the trachea’s direct contact with hemolymph, allowing it to actively acquire lipids, while the midgut is covered by visceral muscle and basement membrane [80, 81].

The mechanism by which VLCFA metabolism regulates tracheogenesis remains unclear, though it may involve the induction of Reactive Oxygen Species (ROS) in the trachea. ROS has been implicated in the activation of HIF-1α/Sima, and VLCFAs is known to induce mitochondrial dysfunction and ROS production [82–84]. Increased ROS levels, through treatments such as paraquat, H_2_O_2_ or bacterial infection, have been shown to induce tracheogenesis in the midgut [42]. Furthermore, manipulating ROS level in the trachea alone can trigger tracheal branching and ISC proliferation [42]. Interestingly, overexpressing *Hnf4* alone in oenocytes could promote tracheogenesis, but does not significantly induce ISC proliferation (Figure S4A). This difference may be attributed to variations in driver strength, with tracheal development occurring prior to ISC proliferation.

Our data indicate that wax esters (WEs) and acylcarnitines (AcCas) are elevated in tumor-bearing flies and mice (Figure 1A and 7N). Notably, in both models we observe an increase in WEs containing very-long-chain fatty acids (VLCFAs), consistent with the elevated expression of *Hnf4* and fatty acid elongases in the KL mouse liver as well as in oenocytes from Yki flies. Functionally, increased AcCas and WEs could contribute to tracheogenesis by supporting local energetic demand. For example, hepatocyte-derived plasma acylcarnitines have been shown to be required for systemic thermogenesis under cold exposure in mice, where peripheral tissues such as brown adipose tissue, skeletal muscle, and heart take up circulating acylcarnitines and oxidize them to fuel metabolism [55]. In addition, wax esters can serve as high-density neutral lipid reserves that are mobilized to meet energetic demands in diverse physiological contexts (e.g., diapause and reproduction in marine organisms). Alternatively, lipid elevation could promote tracheogenesis by increasing reactive oxygen species (ROS), which has been implicated as a driver of tracheal growth . WEs can be hydrolyzed into fatty acids and fatty alcohols [42, 85]. VLCFAs released from WEs may undergo peroxisomal oxidation, generating H₂O₂ as a byproduct, which can stimulate tracheogenesis [42]. In parallel, fatty acids can be transported to mitochondria for β-oxidation, and high fatty-acid flux through mitochondrial respiration can increase electron leak from the ETC and elevate ROS production, potentially contributing to tracheogenesis [86]. These studies also leave open the possibility that metabolites derived from these lipid species act as signaling molecules that promote tracheogenesis.

Additionally, increased AcCas and WEs could contribute to tracheogenesis by supporting local energetic demand. For example, hepatocyte-derived plasma acylcarnitines have been shown to be required for systemic thermogenesis under cold exposure in mice, where peripheral tissues such as brown adipose tissue, skeletal muscle, and heart take up circulating acylcarnitines and oxidize them to fuel metabolism [55]. In addition, WEs can serve as high-density neutral lipid reserves that are mobilized to meet energetic demands in diverse physiological contexts (e.g., diapause and reproduction in marine organisms). These studies also leave open the possibility that metabolites derived from these lipid species act as signaling molecules that promote tracheagenesis.

We observed that VEGF-A stimulates lipid metabolism gene expression in human hepatocytes, and lung tumor-bearing mice showed increased expression of *Hnf4α* and *Elovl7* in peripheral hepatocytes. This work provides insights into tumor-host interactions, challenging the conventional view that tumor-associated metabolic changes are merely passive processes and instead demonstrating their role in promoting tumor growth.

Specifically, inhibiting *Hnf4* and *mElo* in oenocytes slows tumor growth, halts tumor-induced cachexia, and extends the lifespan of flies (Figure 1 and Figure S1). Yki activation increased levels of AcCas, PCs, and TAGs in the hemolymph, and inhibiting *Hnf4* in oenocytes reversed these lipid changes induced by Yki (Figure 4A). This study identifies the TORC1-Hnf4-Elovl7 axis in the oenocytes and tumor-induced lipid metabolism as potential therapeutic targets for managing tumor growth or as biomarkers for early tumor detection. The current data do not exclude the possibility that changes in other Hnf4-expressing tissues contribute to systemic phenotypes, such as bloating, ovary size and lifespan modulation. However, given that oenocyte Hnf4–mElo axis has prominent role in promoting intestinal tumorigenesis, the tumor-induced wasting phenotypes are considered secondary outcomes. Future experiments are needed to investigate how altered Hnf4 activity in muscle or fat body may influence systemic wasting in the Yki tumor model.

Hnf4 is a master regulator of multiple metabolic pathways in *Drosophila* and inhibiting it could have broad effects on the host. For instance, knocking down *Hnf4* in non-tumor-bearing flies led to increased steatosis (Figures S1L). These wide-ranging effects may limit the efficacy of Hnf4 inhibition as an anti-tumor strategy. In contrast, mElo, which is more specifically involved in lipid elongation, presents a more targeted inhibition with lower toxicity and a greater extension of lifespan (Figures 1M, S1O). Our study also suggests a potential role for elongases in tumor-induced angiogenesis, as overexpression of *ELOVL7* has been linked to prostate cancer and is required for cancer growth [87]. While *mElo* is exclusively expressed in oenocytes in flies (Figure 1J), its ortholog, *ELOVL7*, is widely expressed in multiple tissues, including the prostate, kidney, brain, small intestine, and thyroid [88]. It is plausible that *ELOVL7* is upregulated in various tissues in cancer patients, indicating additional tissue targets for anti-angiogenic therapies.

The regulation of *mElo* by Hnf4 seems to be prominent and specific compared to other transcriptional targets under normal physiological conditions (Figure 1I and S1G). It further suggests that the tumor-associated metabolic milieu may reshape Hnf4-driven transcription, potentially by altering (i) cofactor interactions and/or (ii) ligand availability, thereby promoting selective activation of downstream targets such as *mElo*. One limitation of the current study is the absence of mechanistic dissection of how Hnf4 selectively induces mElo expression, largely due to technical challenges associated with obtaining sufficient quantities of purified oenocyte tissue for downstream analyses. Supporting the idea that Hnf4 selectively activates *mElo* under certain conditions, previous studies demonstrate that HNF4*α* transcriptional output is context-dependent and influenced by partner proteins: for instance, HNF4*α* can cooperate with ChREBP (carbohydrate response element-binding protein) to stimulate lipogenic gene *Fasn* during feeding [68, 70], whereas under fasting conditions it can interact with *Foxo1* to enhance expression of gluconeogenic genes such as *G6Pase* [89]. Moreover, HNF4*α* activity can be modulated by long-chain fatty acids [46], raising the possibility that changes in lipid availability in tumor-bearing flies (e.g., due to TAG mobilization) may further influence Hnf4 activity and target selection. Collectively, these findings support a model in which tumor-induced shifts in Hnf4 cofactors and/or lipid ligands redirect Hnf4 transcriptional program in Yki oenocytes, leading to robust *mElo* induction.

In summary, our data reveal a paradigm in which growing tumors actively acquire substrates for tracheogenesis by secreting a PDGF/VEGF-like factor that activates the TORC1-Hnf4 axis in distal oenocytes, inducing the expression of *mElo*. This promotes the synthesis of VLCFAs, which are transported to the trachea to further support tumor growth (Figure 7O).

## Methods

The research described here complies with all relevant ethical regulations.

### Drosophila stocks and husbandry

All experiments were conducted using female flies due to their more consistent bloating phenotype and larger oenocyte, which facilitated dissection. For temperature-sensitive experiments, fly crosses were set up and maintained under a 12:12 hour light:dark cycle at 18°C. Crosses were discarded after 5 days to maintain consistent population density. Following eclosion, adult flies were collected and kept at 18°C for 2–3 days. GAL80^TS^ driver activation was achieved by incubating the flies at 29°C for the durations specified in each experiment. For experiments involving the GeneSwitch driver, fly crosses were maintained at 25°C under a 12:12 hour light:dark cycle and discarded after 3 days. Adult progenies were collected and fed on food containing 200 µM RU486 (Mifepristone, Cayman Chemicals, 10006317), prepared by dissolving the compound in 95% ethanol before adding it to standard fly food. Flies were fed on RU486-containing food for 7 days. Esg^ts^ (*esg-Gal4, UAS-GFP, tub-GAL80^TS^*), *Esg-LexA* and *LexAop-yki^3SA^-GFP, Attp40 UAS-Luciferase* are from the Perrimon lab stock collection.

The following strains were obtained from the Bloomington *Drosophila* Stock Center (BL), Vienna *Drosophila* Resource Center (V) and flyORF (F): *UAS-Hnf4 RNAi* (BL# 29375), *UAS-mElo RNAi* (BL# 44510), *UAS-yki^3SA^* (BL#28817), *UAS-Pvf1-3xHA* (F002862), *UAS-PvR[A]* (BL#58496), *UAS-Pvf1 RNAi* (V#102699), *UAS-PvR RNAi* (V#43459), *UAS-Luciferase RNAi* (BL#31603), Oe^ts^ (*Desat1-Gal4, tubP-Gal80*) (BL# 65406), *UAS-LpR1 RNAi* (BL# 50737), *UAS-LpR2 RNAi* (BL# 54461).

*UAS-Hnf4* is a gift from Dr. Carl Thummel [46]; *UAS-TSC1, TSC2* is a gift from Dr. Christen K Mirth [90] and UAS-PRAS40 was a gift from Dr. Aurelio Teleman [91], Btl^TS^ (*Btl-Gal4, UAS-srcGFP*) and *QF6 >QUAS-mtdTomato* was a kind gift from Dr. Chrysoula Pitsouli [42]. PromE^GS^(*PromE-GeneSwitch-Gal4*) was a gift from Dr. Heinrich Jasper [92]. *UAS-CG18609* and *mFas-Gal4* were gifts from Dr. Henry Chung [93, 94]. *UAS-PRAS40* is a gift from Dr. Aurelio A Teleman [74].

Hnf4-127D01 was generated through CRISPR-mediated tagging of endogenous Hnf4 with 127D01-tag as described previously [47]. Briefly, we cloned the sequence 1 kb upstream and 1 kb downstream of the stop codon into the donor vector pScarlessHD-C-3x127D01-DsRed (Addgene#171578). A sgRNA plasmid pCFD3-Hnf4-sgRNA that target the seed sequence (CCAGAGACTGGTTACTAGAAGAA) near the stop codon of Hnf4 was injected into yw;nos-cas9/CyO embryos together with the donor plasmid. Positive transformants with red eye fluorescent eyes were outcrossed and successful Kis were confirmed by PCR (F: GTACAACCGGAGCGAGGGTA; R: CAAAATGGTTCAAGGCATAACA) and sequencing.

Subsequently, 3xP3dsRed was excised using piggyBac transposase (BL#8285).

### Untargeted lipidomic and metabolomic analysis

Lipidomics and metabolomics of fly samples was performed following the Folch method (Folch et al., 1957). 15 adult flies, 20ul of mice serum and hemolymph from 200-300 adult flies were used for lipidomics and metabolomics, with 3-4 biological replicates. Solid samples were homogenized in a 1 mL Dounce homogenizer (TIGHT) on ice with 0.6 mL chloroform and 0.3 mL methanol (2:1 chloroform-methanol mixture). Homogenized samples were transferred to a 15 mL glass tube with a Teflon cap, and an additional 0.6 mL chloroform and 0.3 mL methanol were added to make the final volume approximately 1.8 mL. Hemolymph samples or conditioned medium was transferred to a 15 mL glass tube containing 2:1 chloroform-methanol mixture. Samples were vortexed briefly and incubated at room temperature on a rotator for 30–45 minutes. Following incubation, 0.2 volumes (0.36 mL) of HPLC-grade deionized water (HPLC dH2O) were added. The mixture was vortexed three times, followed by centrifugation at 1000 × g for 10 minutes at 4°.

The upper aqueous phase, containing polar metabolites, was carefully collected using a glass pipette, while avoiding the middle phase. Metabolites were transferred to microcentrifuge tubes, evaporated under vacuum using a SpeedVac rotary evaporator overnight, and stored at -80°C. Polar metabolites were resuspended in 20 μL of HPLC-grade water and then injected and analyzed using a hybrid 6500 QTRAP triple quadrupole mass spectrometer (AB/SCIEX) connected to a Prominence UFLC HPLC system (Shimadzu). The lower phase, containing non-polar lipids, was transferred to a clean 1.85 mL glass test tube using a glass pipette. The phase was dried under nitrogen gas for approximately 30 minutes, and dried lipids were stored at -80°C to prevent oxidation. Non-polar lipids were resuspended in 35 μL of HPLC-grade 50% methanol/50% isopropyl alcohol (IPA) and then injected and analyzed using untargeted high-resolution LC-MS/MS on a Thermo QExactive Plus Orbitrap mass spectrometer. LipidSearch version 4.2 (Thermo Scientific) was used to identify lipid molecules, with the quantification of ion intensity by measuring the area size of identified peaks.

The full names of the lipid species can be found in Supplementary Data 4. For lipidomic quantification using whole fly bodies, the same number of flies was used for lipid extraction. Lipid species were normalized to the total reads in each sample. For hemolymph lipidomic analysis, lipid species were normalized to both the volume of hemolymph extracted and the total reads in each sample.

### RNA extraction Quantitative RT-PCR

Adult female fly oenocytes were dissected in cold 1xPBS before RNA extraction. For oenocyte dissection, the fat body was removed through liposuction and then oenocytes were removed from the cuticle using a fine-tip glass needle. Tissue lysis, RNA extraction, and cDNA synthesis were performed using Cells-to-CT Kit (Thermo Scientific, #44-029-53). For female adult whole body mRNA extraction, four flies per biological replicate were collected. Total RNA was extracted using Direct-zol RNA MicroPrep kit (Zymo Research, #R2060). mRNA was extracted from HepG2 cells using Directo-zol RNA MicroPrep kit (Zymo Research, #R2060). 30-50mg of mice livers were dissected and total RNA was extracted using Qiagen RNeasy Mini kit (Qiagen #74104). All the cDNAs were synthesized using the iScript cDNA synthesis kit (Bio-Rad, #1708896) in a 40ul reaction mixture using 1000ng total RNA. Quantitative real-time RT-PCR (RT-qPCR) assays were performed using iQ SYBR Green Supermix (Bio-Rad, #1708880) on a CFX96 Real-Time PCR Detection system (Bio-Rad). Two to four independent biological replicates were performed with two technical replicates. The mRNA abundance of each gene was normalized to the expression of *RpL32* or *αTub84B* for fly samples, Gapdh for mouse samples and ACTB for HepG2 samples. Primer sequences are listed in the Supplementary Data 5.

### Fly imaging

*Drosophila* ovaries were dissected in cold PBS. ovaries were imaged immediately in cold PBS using a ZEISS Axiozoom V16 fluorescence microscope. Flies were anesthetized on CO_2_ pad and imaged using ZEISS Axiozoom V16 fluorescence microscope. The size of the ovary and abdomen is measured using Fiji (version: 2.14.0/1.54f).

### Oil Red O staining

The method was as previously described [29]. Flies were dissected in PBS and fat bodies were removed with liposuction. Specimens were fixed in 4% paraformaldehyde for 20 minutes. Specimens were rinsed twice with distilled water. Oil Red O (O0625-, Sigma-Aldrich) was dissolved in isopropanol to make 0.1% Oil Red O stock. A working solution was prepared by combining 3 mL of 0.1% Oil Red O in isopropanol with 2 mL of distilled water. The solution was freshly prepared and filtered through a 0.2-µm syringe filter before use. Specimens were incubated for 25 minutes in Oil Red O stain, rinsed briefly with PBS and mounted in Vector Shield for imaging. Oil Red O images were taken using Olympus VS200 Slide Scanner, equipped with color camera (2448x2048 pixels, 3.45μm).

### Immunostaining and image analysis

For immunostaining using the nanobodies, fly abdomens were dissected in PBS at room temperature with fat bodies removed and fixed in 4% formaldehyde in PBS for 30 minutes. Specimens were washed three times in PBST (0.1% Triton-X-100), 5 minutes each. The specimens were then incubated in 5% NGS (normal goat serum in PBST) for 1 hour.

Nb127D01-ALFA, 0.2mg/ml was used at a dilution of 1:500, incubating the samples for 1 hour and a half with mild shaking. Followed by 0.1% PBST washes, 5 minutes each wash. AlexaFluor 594 anti-Alpaca (Jackson Immuno Research, 128-585-232) was diluted 1:400 in PBS-TX + 5% NGS and incubate with the specimens at room temperature with mild shaking for 1 hours. After washes and DAPI staining, tissues were mounted in Vectashield and imaged with Olympus VS200 Slide Scanner or confocal microscope Zeiss Axio Observer Z1.

For regular immunostainings, fly tissues were dissected in PBS at room temperature.

Tissues were fixed in 4% paraformaldehyde in PBS for 30 minutes. Specimens were washed three times in 0.1% PBST and incubated in 5% NDS solution for 1 hour. Tissues were then incubated with primary antibodies diluted in PBST and blocking solution overnight at 4°.

Then the tissues were washed three times with PBST at room temperature. Secondary antibodies were applied for 1 hour. Excess antibodies were washed in PBST, and tissues were incubated with DAPI (Invitrogen, D1306) dilute in PBST for 5 minutes. Tissues were then mounted in Vectashield (Vector Laboratories Inc, H-1000). The following primary antibodies were used: rabbit-anti-pH3 (1:500, Cell Signaling, 9701), rabbit-anti-p4E-BP (1:800, Cell Signaling, 2855S). Secondary antibodies against mouse, rabbit, chicken or alpaca conjugated to Alexa Fluor 488 and 594 (Jackson Immuno Research) were used at 1:800 for anti-p-4EBP, 1:1000 for anti-pH3 and 1:400 for Nb127D01-ALFA. For immunostaining in HepG2 cells, cells were washed twice with PBS and fixed in 4% paraformaldehyde for 25 min, followed by a PBS wash. Samples were permeabilized with 0.5% Triton X-100 in PBS for 10 min and washed with PBS. Non-specific binding was blocked in 1% BSA in PBS for 1 h at room temperature. Cells were then incubated with anti–HNF4α rabbit monoclonal antibody (C11F12, Cell Signaling Technology, #3113) diluted 1:1000 in PBS at 4 °C overnight. After washing twice with 0.05% Triton X-100 in PBS, samples were incubated with Alexa Fluor 594–conjugated secondary antibody (1:500 in PBS) at room temperature. Cells were washed three times with 0.05% Triton X-100 in PBS and once with PBS, then mounted using SlowFade® Gold antifade reagent with DAPI (Invitrogen, S36939) and sealed with a coverslip.

Quantification of nucleus Hnf4 intensity was conducted using ImageJ. Background subtraction was performed using the rolling ball radius at 50 pixels. We apply threshold to the DAPI channel and using analyze particles to select nucleus as Region Of Interest (ROI), size: 10-2000; circularity: 0.1-1. Mean gray value was measured for each ROI.

### Lifespan analysis

For survival analysis, flies were collected within 24 hours post-eclosion (20 females and 3 males per vial) and maintained on standard laboratory fly food at 18°C, with 60% humidity, and a 12-hour light/dark cycle. After two days, mature and mated females were transferred to 29°C to induce gene expression. To maintain vial cleanliness, flies were transferred to fresh vials every two days, and the number of dead flies was recorded daily.

### snRNAseq analysis

Data and plots from Figure 1 were generated from the Fly Cell Atlas [95], generated from the 10X platform, “Stringent All” dataset. Other snRNAseq dataset was obtained and analyzed as described in [50], where a resolution of 0.4 was chosen as the clustering parameter.

For the Seurat Module Score analysis, Seurat AddModuleScore function was used to calculate the average expression of a group of genes in Very Long Chain Fatty Acid Synthesis pathway in each cell (for details of the list, see Supplementary Data 6), then subtracted by a randomly picked gene set expression.

### Rapamycin feeding

Rapamycin was dissolved in DMSO to make 2mM rapamycin stock. Rapamycin stock was further diluted to 200μM in ethanol and evenly distributed on to food surface. A small piece of Kimwipe was used to remove excess liquids on the food. DMSO dissolved in ethanol was used as control. Flies were transferred to new food every two days. Flies were fed for 6-7 days before dissection.

### Dextran injection

Using a fine glass needle, ∼100nl of 50 mg/ml 10,000 MW TR-dextran (Thermo Fisher Scientific, D-1817) diluted in water was injected in the abdomen of adult females. 6 hours after injection, abdomen and thoraces were separated and whole abdomen were fixed in 4% paraformaldehyde for 30 minutes with mild shaking. After fixation, midguts were dissected and washed briefly with PBS and mounted in Vectashield. Midguts were imaged using a single point laser scanning confocal, Olympus IX83 to measure the trachea leakage. A Z-stack was taken, and maximum projections were used to generate final image for analysis.

### Trachea imaging

Generating fly strains that incorporate both a trachea reporter and other genetic modifications is challenging. Therefore, the following method, as described in [42], was used to quantify the trachea. Flies were dissected in cold PBS, and intestines were carefully extracted to maintain their original structure. Tissues were then fixed in 4% paraformaldehyde for 20 minutes, mounted in Vectashield, and imaged within 3 days. Whole intestine images were captured using an Olympus VS200 Slide Scanner equipped with a color camera. Only the upper layer of the tissue, in contact with the coverslip, was analyzed.

### Trachea analysis

To automate trachea measurement, we used the previously developed software AutoTube [96]. Only the topmost Z-stack, which provided the clearest image of the trachea, was analyzed. Images corresponding to the R2-R4 regions of the gut were cropped, avoiding areas with the major trunk of the trachea to prevent skewing the results. Images were thresholded on a white background and analyzed with AutoTube using the following settings:

- Input Color Channel: Green
- Intensity adjustment: Autocontrast
- Illumination correction: Illumination size set to 51
- Noise correction: BM3D
- Tube detection: Finer tubes with a tube size of 3
- Small regions removed: 0.01% of Ima
- Hole filling: Hole size set to 1
- Threshold type: MultiOtsu
- Short ramifications removed: Spur length set to 15
- Spatial branch merging: Spatial distance set to 10

The analysis provided results for "Number of Branches (μm)," "Total Tube Area (μm)," and "Total Skeleton Length (μm)," which were normalized to the corresponding intestinal area and were multiplied by an arbitrary number 1000000. Results from the R2-R4 regions were averaged for each gut.

### Animal care and tumor induction

*Kras^G12D^/+;Lkb1^f/f^* mice, previously described [7, 97], were backcrossed to the FVB strain. Male mice were maintained on a 12-hour light/dark cycle at an ambient temperature of 22°C, with ad libitum access to rodent chow (PicoLab Rodent 5053; Lab Diet, 3.43 kcal/g) and drinking water. Tumors were induced in adult male mice (12 to 20 weeks old) via intranasal administration of 75 µL PBS containing 2.5 × 10⁷ pfu of adenovirus SMV-CRE (Ad5CMV-Cre) purchased from the University of Iowa Gene Transfer Vector Core (Iowa City, IA). The endpoint cutoff for these experiments was 25% weight loss. These experiments used an orthotropic lung cancer model, and thus there was no specific tumor size endpoint. The ages of the mice ranges from 7-38 weeks. All animal experiments were conducted with the approval of the Institutional Animal Care and Use Committee (IACUC) at Weill Cornell Medicine (WCM) under protocol number 2013-0116.

### Mice tissue collection

Male mice were euthanized with CO_2_, followed by blood collection via cardiac puncture. Liver, gonadal adipose tissue, and skeletal muscles were dissected and flash-frozen in liquid nitrogen and stored at -80°. The control mice were a mix of fasted and fed mice. “Fasted” mice were fasted for 12 hours and “fed” mice were fasted for 12 hours and then allowed free access to food for 4 hours before tissue collection.

### VEGFA incubation

HepG2 cells (obtained from ATCC) were cultured in HyClone Dulbecco’s Modified Eagle Medium (DMEM) with high glucose (SH30022.01): with L-glutamine; without sodium pyruvate. Additional 50U penicillin/ml and 50ug streptomycin/ml were added. HepG2 cells were seeded at density of 5 x 10^5^ in 12-well-plate (VWR, 29442-040), treated or not with 50ng/ml Recombinant Human VEGF 165 (PeproTech, 100-20-2UG) for 24 hours. Total cellular RNA was isolated using the Trizol reagent and extracted through Direct-zol RNA MicroPrep kit (Zymo Research, #R2060).

### Statistics & Reproducibility

Sample size was determined according to previous published studies [12, 21, 71, 98]. No data were excluded from the analyses, and the experiments were randomized and no blinding was done.

## Data Availability

Data generated and used in this study are described within the Article, Supplementary Data files, Supplementary Information, and Source Data file. Source data is provided in this paper.

## Code Availability

See code for generating plots for snRNAseq, and methods for analyzing lipidomic data here https://github.com/carriehuang1993/snRNAseq-analysis.git [99].

## Acknowledgments

We thank Dr. John M. Asara at the Beth Israel Deaconess Medical Center for his help on lipidomic analysis. We thank the assistance provided by the Microscopy Resources on the North Quad (MicRoN) core and we thank Dr. Simon F. Norrelykke at the Image Analysis Collaboratory at Harvard Medical School for advice on AutoTube. We also thank Dr. Ah-Ram Kim, Dr. Sun Jin Moon at Harvard Medical School, Dr. Hua Bai at Iowa State University, Dr. Eileen White and Dr. Maria Gomez- Jenkins at Rutgers University for scientific discussion and insights. We thank Dr. Ju Xu at Chinese Academy of Sciences for the help generating Hnf4-127D01 fly strain. We thank Dr. Ying Liu at Harvard Medical School for sharing fly lines used in this study. We also thank Christians Villalta for the help on fly dextran injection.

This article is subject to HHMI’s Open Access to Publications policy. HHMI lab heads have previously granted a nonexclusive CC BY 4.0 license to the public and a sublicensable license to HHMI in their research articles. Pursuant to those licenses, the author-accepted manuscript of this article can be made freely available under a CC BY 4.0 license immediately upon publication.

## Author Contributions Statement

Conceptualization: KH, TM, MG, NP; Methods, Data Acquisition, Resources, Visualization and Imaging: KH, TM, YC; Mice sample preparation: ED, JS; Data Acquisition: KW, Bioinformatic Analysis: MH, YH, SJM; JMA performed Independent replication of experiments at Harvard Medical School: KH; Writing – original draft: KH, Writing – review, editing & revising: KH, TM, NP.

---

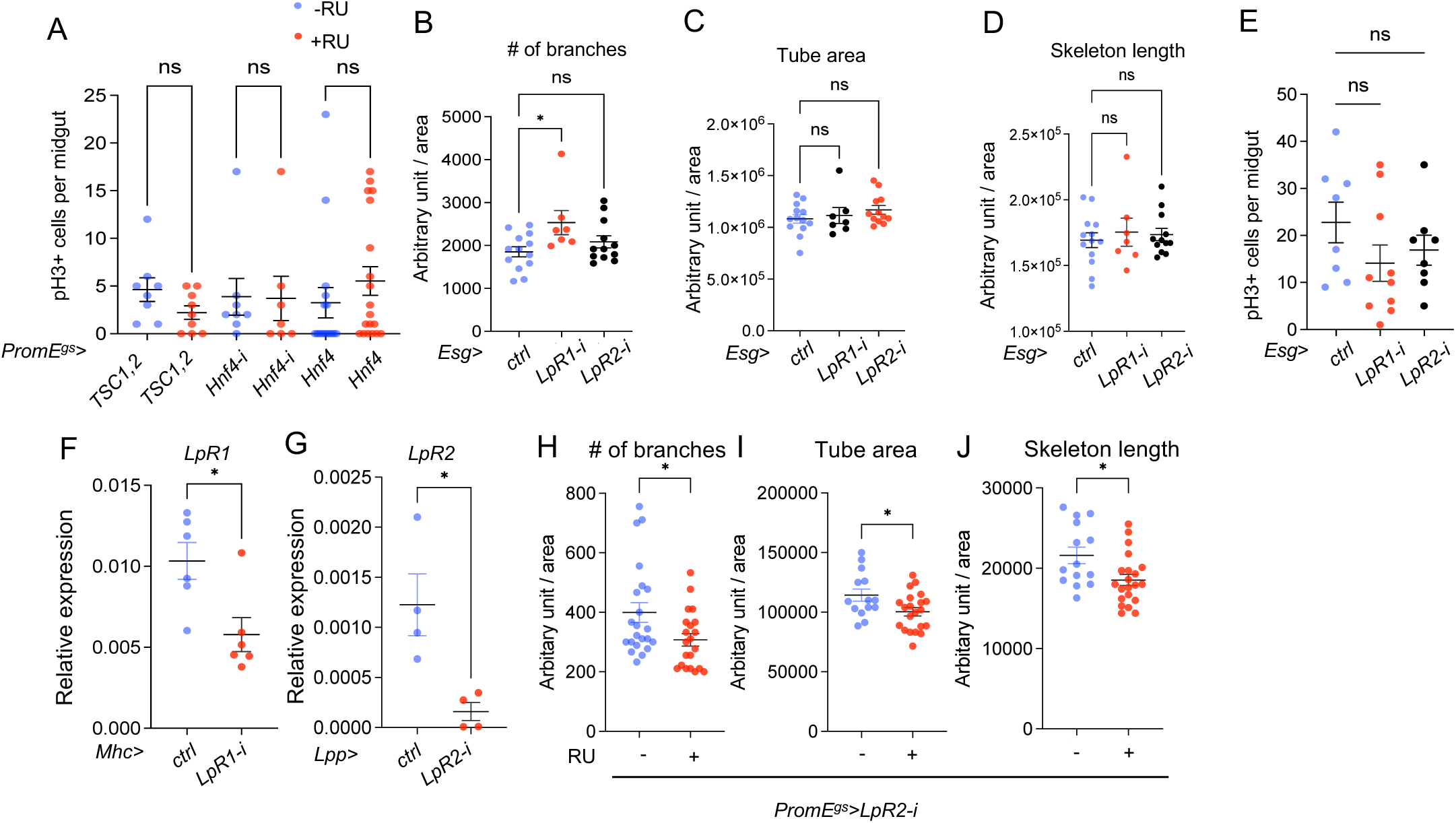

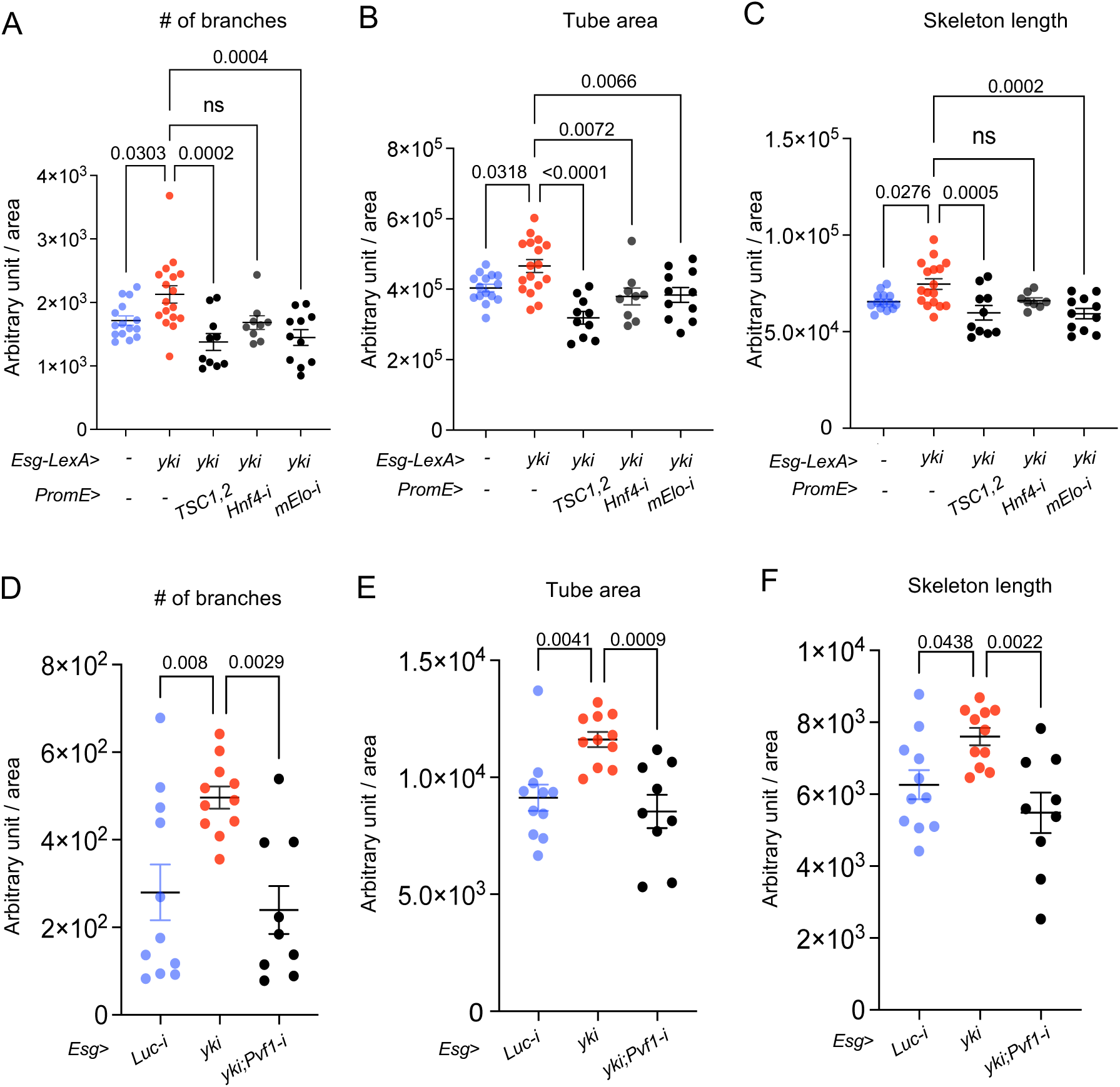

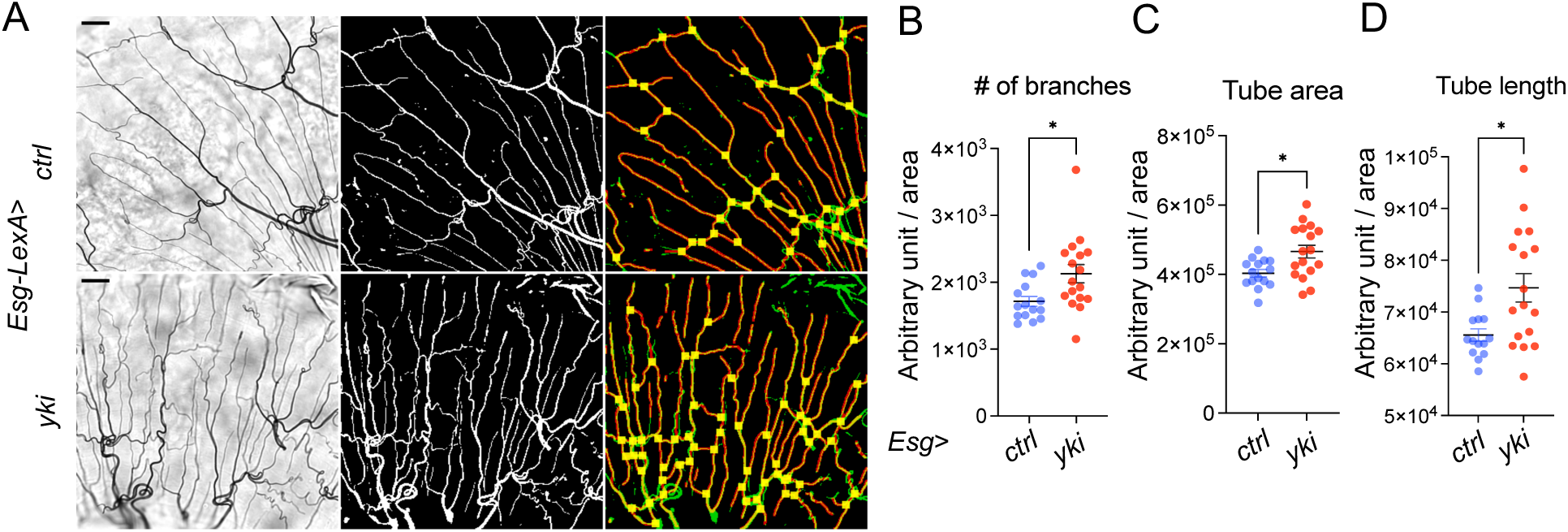

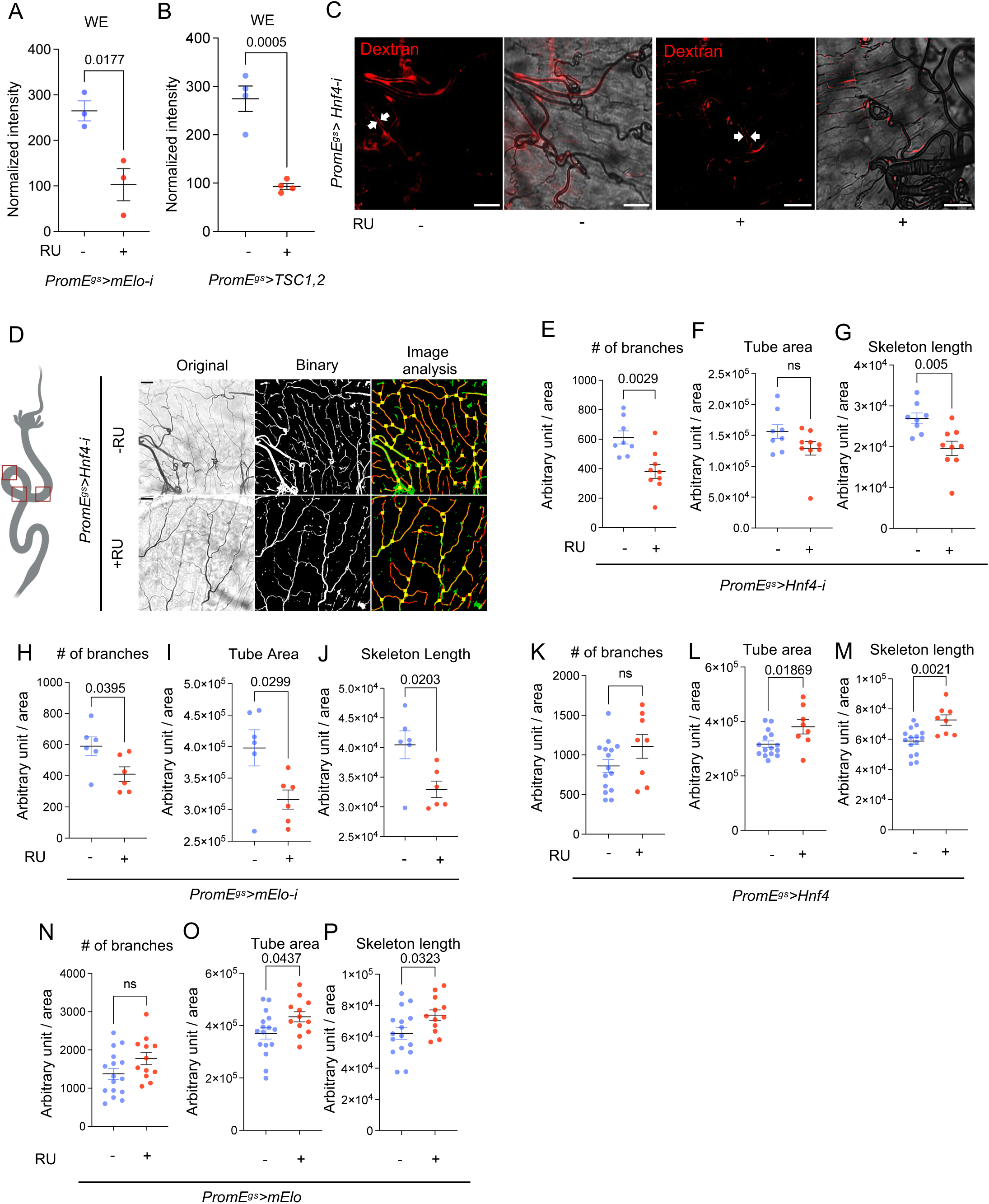

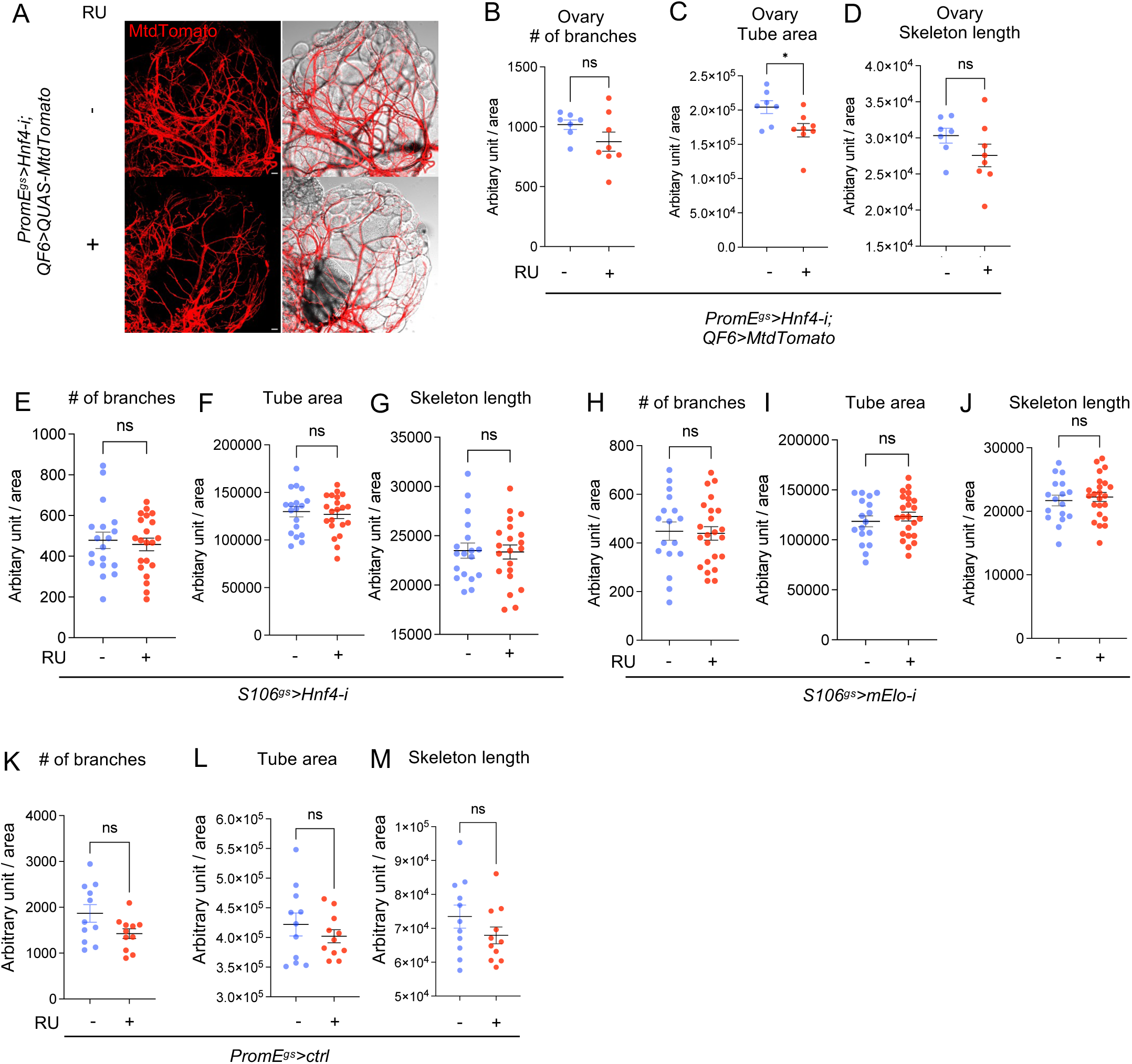

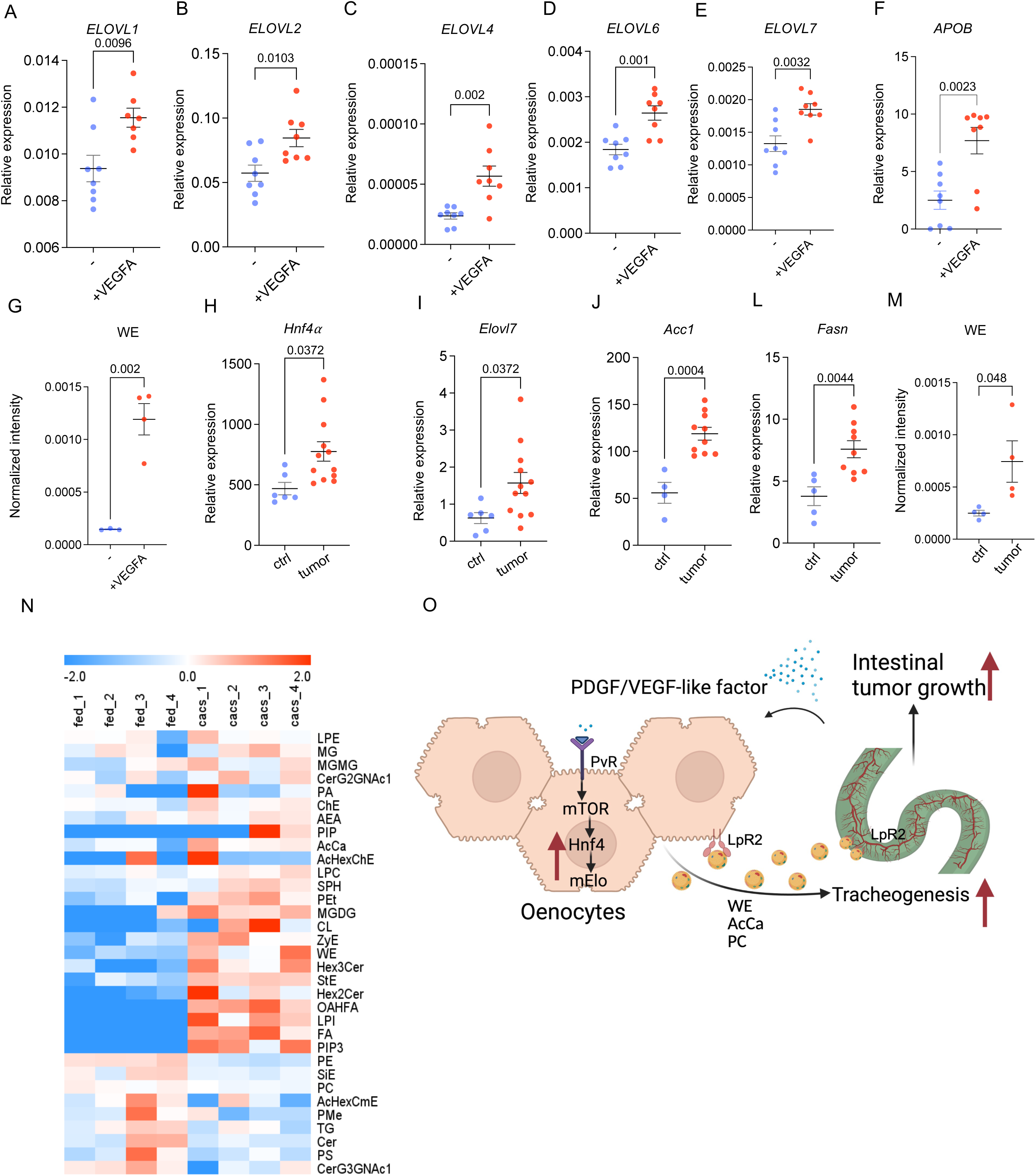

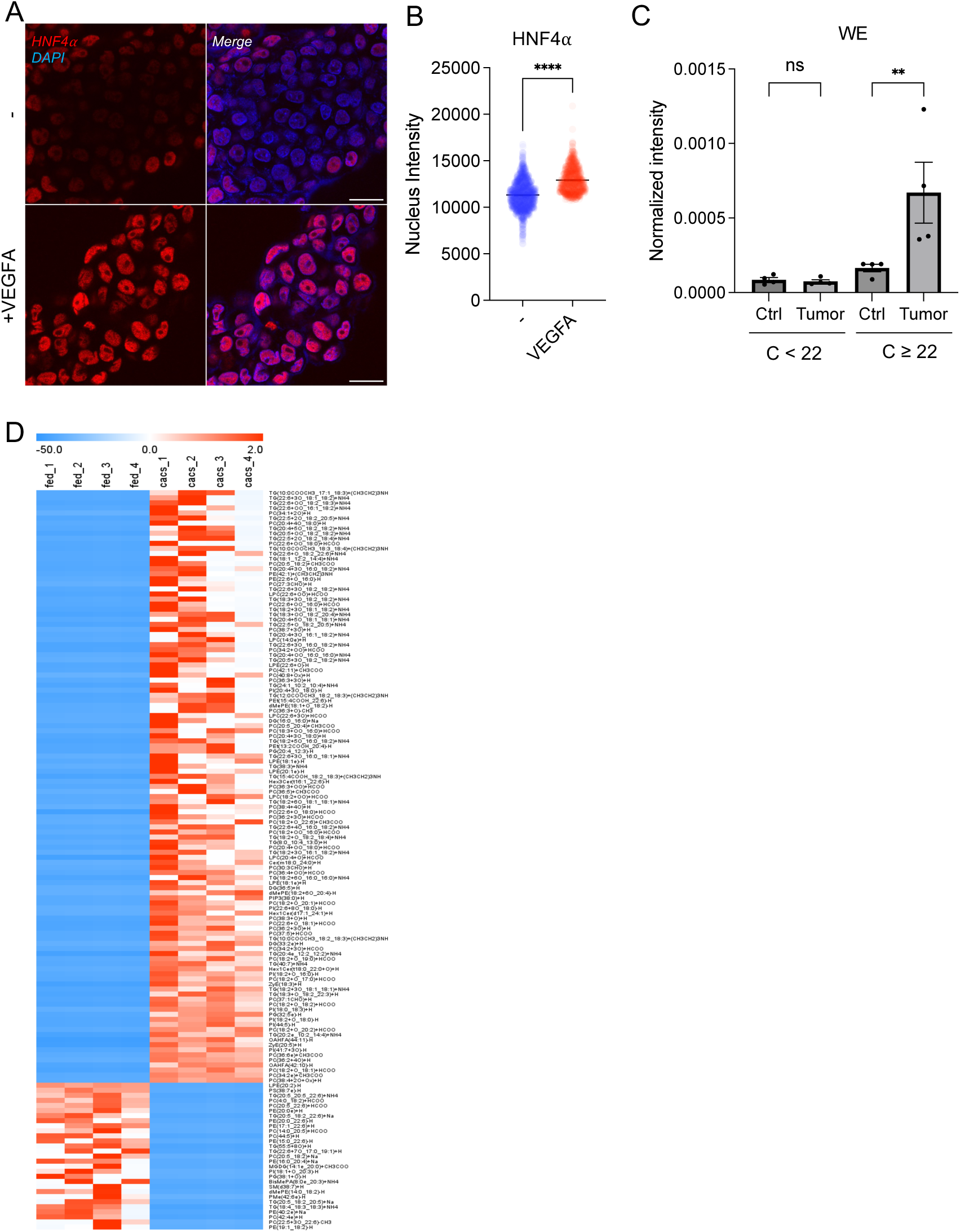

