## Supplementary Figures for "Lipid metabolism of hepatocyte-like cells supports intestinal tumor growth in *Drosophila*"

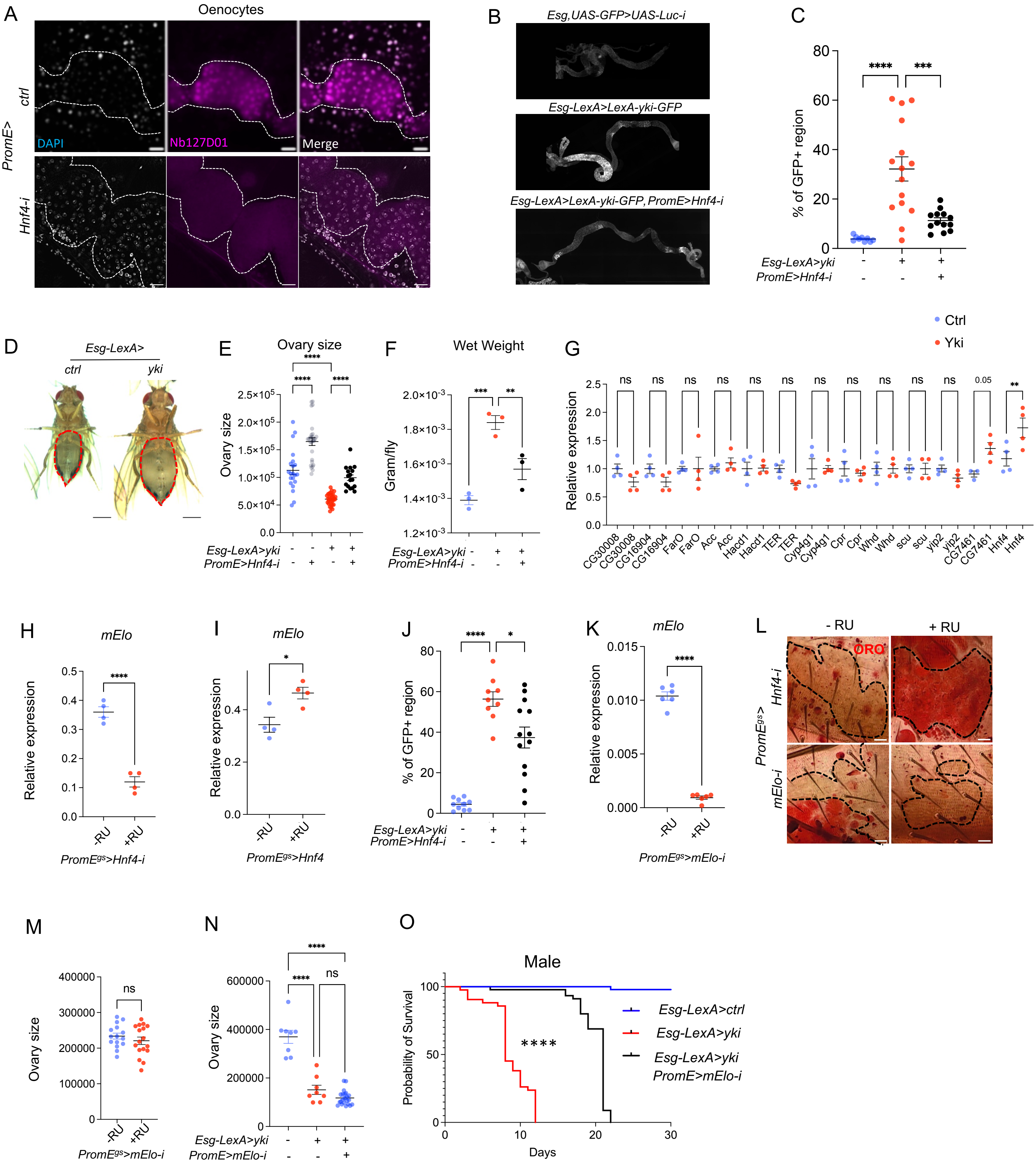
A
Oenocytes
PromE>
ctrl
Hnf4-i
DAPI
Nb127D01
Merge
B
Esg,UAS-GFP>UAS-Luc-i
Esg-LexA>LexA-yki-GFP
Esg-LexA>LexA-yki-GFP, PromE>Hnf4-i
C
% of GFP+ region
Esg-LexA>yki
PromE>Hnf4-i
D
Esg-LexA>
ctrl
yki
E
Ovary size
F
Wet Weight
Gram/fly
G
Ctrl
Yki
Relative expression
CG30008
CG16904
FarO
Acc
Hacd1
TER
Cyp4g1
Cpr
Whd
scu
yip2
CG7461
Hnf4
H
mElo
PromE^gs>Hnf4-i
I
mElo
PromE^gs>Hnf4
J
% of GFP+ region
K
mElo
PromE^gs>mElo-i
L
- RU
+ RU
ORO
PromE^gs>
Hnf4-i
mElo-i
M
Ovary size
PromE^gs>mElo-i
N
Ovary size
Esg-LexA>yki
PromE>mElo-i
O
Male
Probability of Survival
Days
Esg-LexA>ctrl
Esg-LexA>yki
Esg-LexA>yki PromE>mElo-i

**Figure 1S: Hnf4 and mElo regulate metabolic and tumor-related phenotypes in Yki flies.**

(A) Immunostaining of Nb127D01 for nuclear-localized, endogenously tagged Hnf4 (Hnf4-127D01) in oenocytes with *Hnf4* knockdown or control flies, with dashed lines enclosing oenocytes. (B-C) Level of GFP positive region in the whole fly midgut in control, oenocyte specific *Hnf4-i* and *Yki* flies with *Hnf4-i*. (D) Abdominal area of control and Yki flies for bloating measurement. (E) Ovary size in control flies, oenocyte specific *Hnf4-i* flies, *Yki* flies and *Yki* flies with oenocyte specific *Hnf4-i*, n = 9 - 15. (F) Wet weight measurement of fly whole body, where flies were weighed first and were dried at 65°C for 5 hours, the second value was subtracted from the first to obtain the wet weight of the fly. N = 3 biological replicates. (G) Expression of VLCFA biosynthesis and β-oxidation genes in control and Yki flies. (H-I) *mElo* expression measured by qPCR in oenocyte specific *Hnf4* knockdown or *Hnf4* overexpression, N = 2 biological replicates from 2 independent experiments. (J) Level of GFP positive region in the whole fly midgut in control, Yki flies, oenocyte specific *Hnf4*-i and *Yki* flies. (K) Confirmation of *mElo* knockdown efficiency via qPCR, N = 2 biological replicates from 2 independent experiments. (L) Oil Red O staining in oenocytes of *Hnf4* knockdown or *mElo* knockdown. N = 8 flies. (M) Ovary size measurement in control and *mElo-i* flies, N = 13-16 flies. (N) Ovary size in control, Yki and Yki with *mElo-i* flies. (O) Lifespan analysis in Yki-expressing flies upon oenocyte-specific *mElo* knockdown in male flies. Data represent mean ± s.e.m.; significance assessed using appropriate statistical tests. Ns is P > 0.05, *P < 0.05, **P < 0.01, ****P < 0.0001.

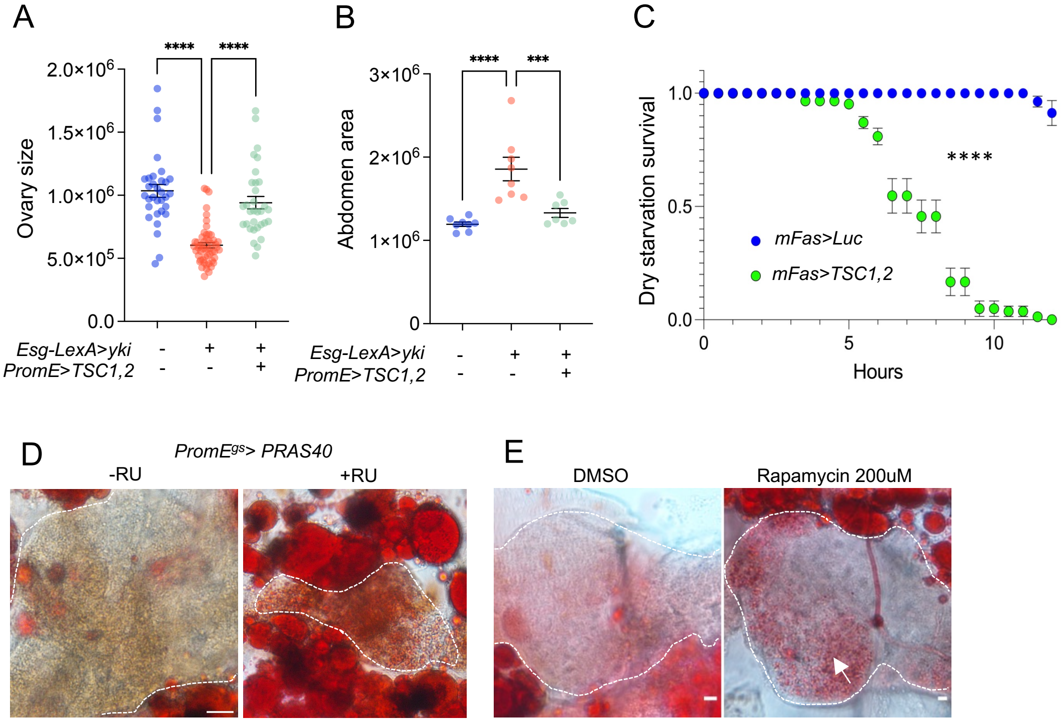


**Figure 2S: The role of oenocyte TORC1 in regulating lifespan, steatosis and Yki wasting phenotype.**

(A-B) Ovary size measurements and bloating are rescued by *TSC1,2* overexpression in Yki flies, N = 15-30 flies. (C) Sensitivity to dry starvation in flies overexpressing *TSC1,2* in oenocytes compared to controls. Adult oenocyte specific driver (*mFas-Gal4*) was used [93]. (D-E) ORO staining indicates steatosis in oenocytes with *PRAS40* overexpression, or in rapamycin treatment. Dashed lines enclose oenocytes in all images. Data represent mean ± s.e.m.; statistical significance determined using Student's t-test or ANOVA as appropriate. Ns is P > 0.05, *P < 0.05, **P < 0.01, ****P < 0.0001.

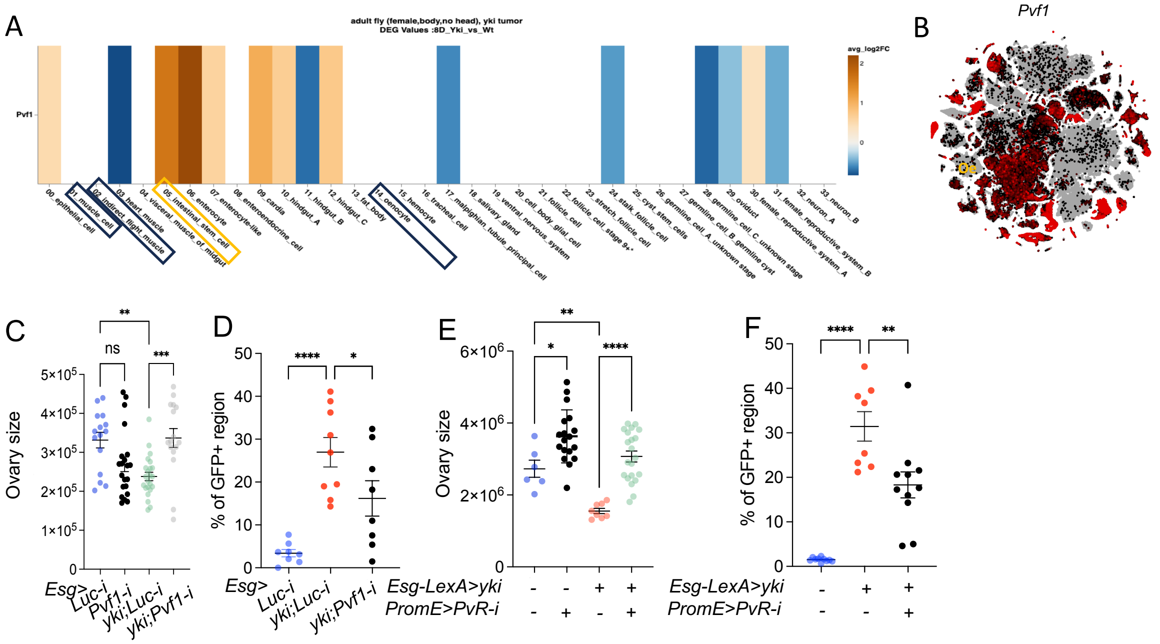

**Figure 3S: Expression of *Pvf1* in fly whole body and oenocytes in Yki flies.**

(A) Heatmap showing average gene expression changes (log2 fold change; Yki vs WT at day 8) across multiple tissues. (B) UMAP from whole body (without head) snRNAseq, shows *Pvf1* expression level, oenocyte cluster is marked as “oe”. (C) Ovary wasting of Yki flies with ISC-specific *Pvf1* knockdown. Dashed lines enclose oenocytes. N = 10-20 biological replicates. (D) Tumor mass (GFP-positive region of the gut) in Yki flies with ISC-specific *Pvf1* knockdown. N = 8-10 biological replicates. (E) Ovary wasting of Yki flies with ISC-specific *Pvf1* knockdown. Dashed lines enclose oenocytes. N = 10-20 biological replicates. (F) Tumor mass (GFP-positive region of the gut) in Yki flies with ISC-specific *Pvf1* knockdown. N = 8-10 biological replicates. Ns is P > 0.05, *P < 0.05, **P < 0.01, ****P < 0.0001.

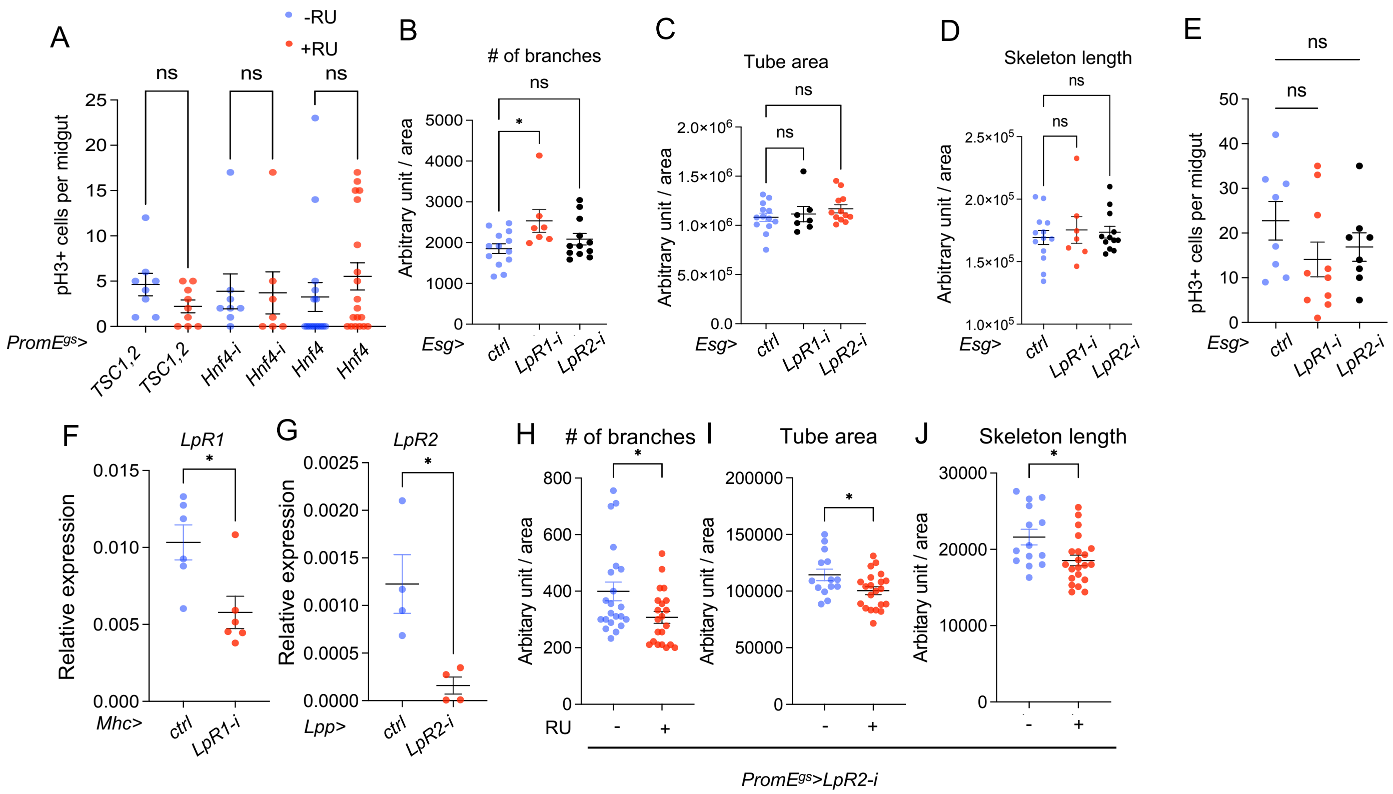


**Figure 4S: Effects of ISC-specific *LpR1* and *LpR2* knockdown on tracheal morphology and ISC proliferation.** (A) Counts of pH3+ cells in the midgut of flies with oenocyte-specific *TSC1*,2 overexpression or *Hnf4* knockdown or overexpression. N = 8-20 flies. (B-D) Trachea parameters, including number of branches, tube area, and skeleton length, in flies with ISC-specific knockdown of *LpR1* and *LpR2* using *Esg* driver, compared to control (*Esg*>*ctrl*). N = 7-11 flies. (E) Counts of pH3+ cells in the midgut of flies with ISC-specific knockdown of *LpR1* and *LpR2*, compared to controls. (F) *LpR1* expression level in fly thoraces with *LpR1* knockdown using temperature sensitive muscle specific driver (Mhc-Gal4) activated at adult stage for 7 days. (G) *LpR2* expression level in fly fat bodies with *LpR2* knockdown using temperature sensitive fat body specific driver (Lpp-Gal4) activated at adult stage for 7 days.  (H-J) Gut tracheal parameters in flies with oenocyte-specific *LpR2* knockdown. N=20-25 flies. Data are presented as mean ± s.e.m.; statistical analysis was performed using Student's t-test or ANOVA as appropriate. Ns is P > 0.05, *P < 0.05, **P < 0.01, ****P < 0.0001.

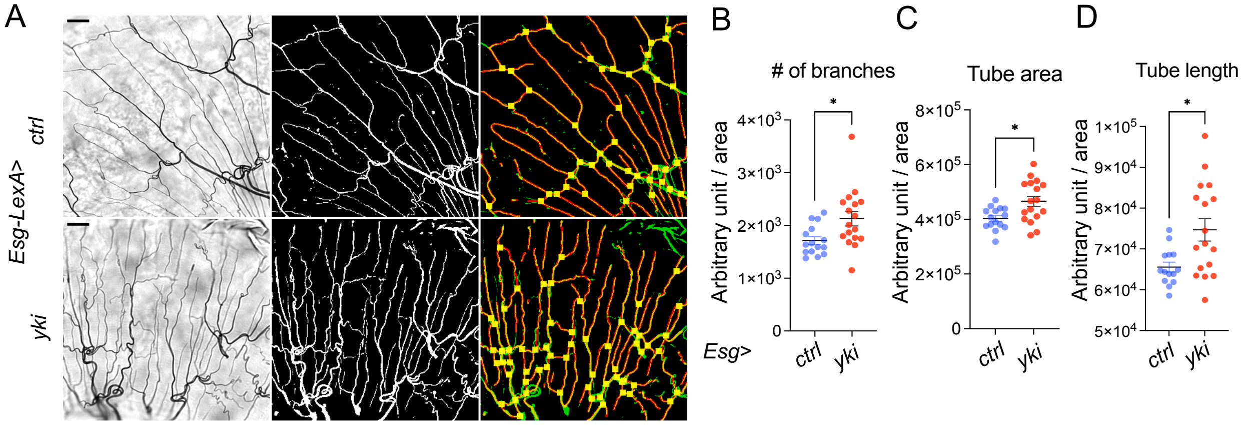


**Figure 5S: Hnf4-mElo axis in oenocytes mediates lipid metabolism and tracheogenesis induced by Yki tumors.**

(A-D) Tracheal morphology parameters, including number of branches, total tube area, total tube length, and skeleton length, in Yki midgut (*Esg*>*yki*), compared to wild-type controls (*Esg*>*ctrl*). N = 16-17. Ns is $P > 0.05$, *$P < 0.05$, **$P < 0.01$, ****$P < 0.0001$.

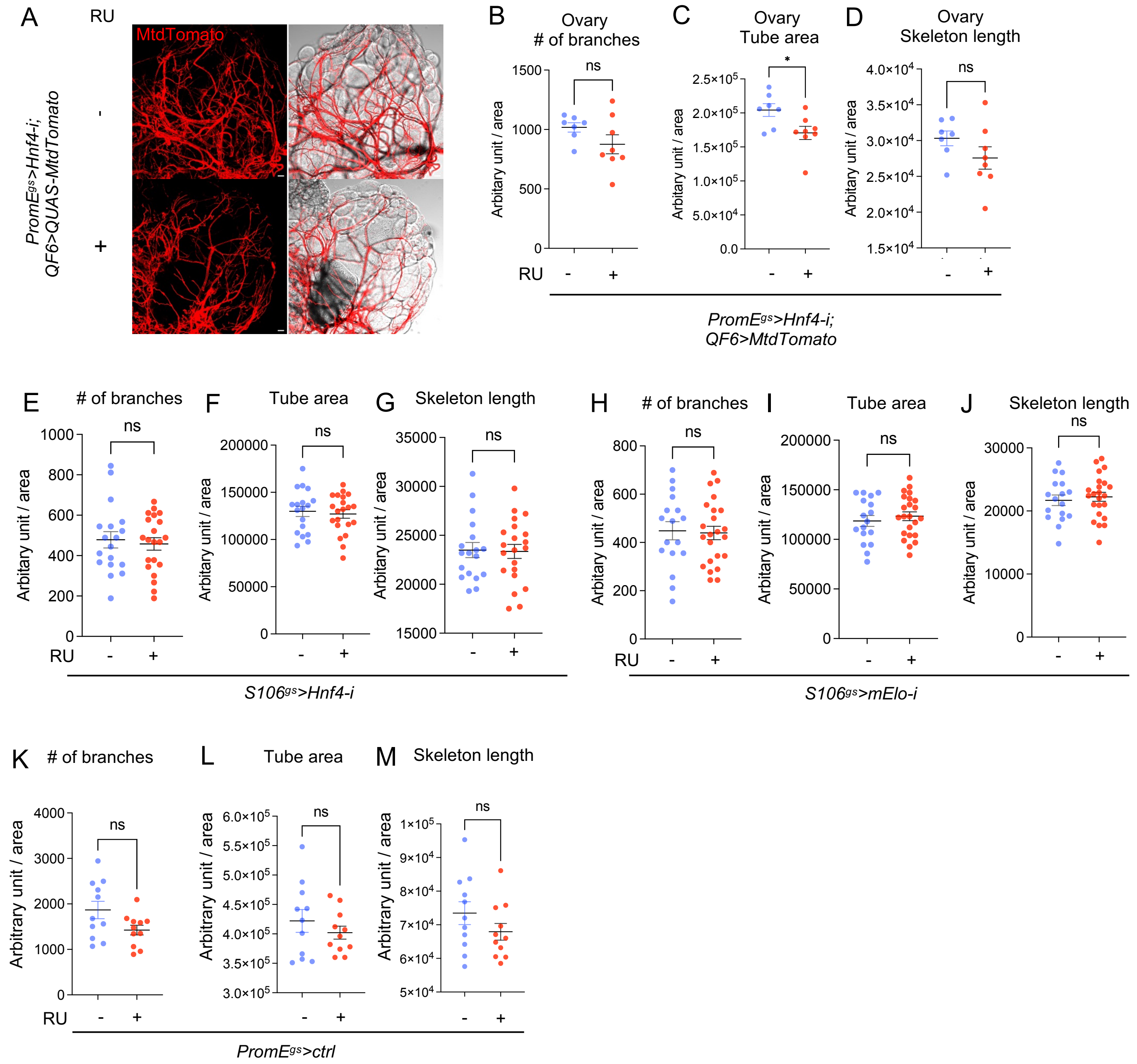


**Figure 6S: Effect of oenocyte Hnf4 on ovary trachea.**

(A-D) Tracheal branch number, tube area and total skeleton length of the ovary in flies with *Hnf4* knockdown in oenocytes. Red (mtdTomato) marks the trachea (QF6>QUAS-mtdTomato). N = 8 flies. (E-G) Tracheal parameters of mid gut in flies with *Hnf4* knockdown in the fat body and gut. (N =18). (H-J) Tracheal parameters of mid gut in flies with *mElo* knockdown in the fat body and gut. (N =18). (K-M) Comparison of gut trachea morphology in control flies (*PromE*$^{gs}$>+) with and without RU feeding. N = 12 flies. Data are presented as mean ± s.e.m.; statistical analysis was performed using Student's t-test or ANOVA as appropriate. Ns is P > 0.05, *P < 0.05, **P < 0.01, ****P < 0.0001.

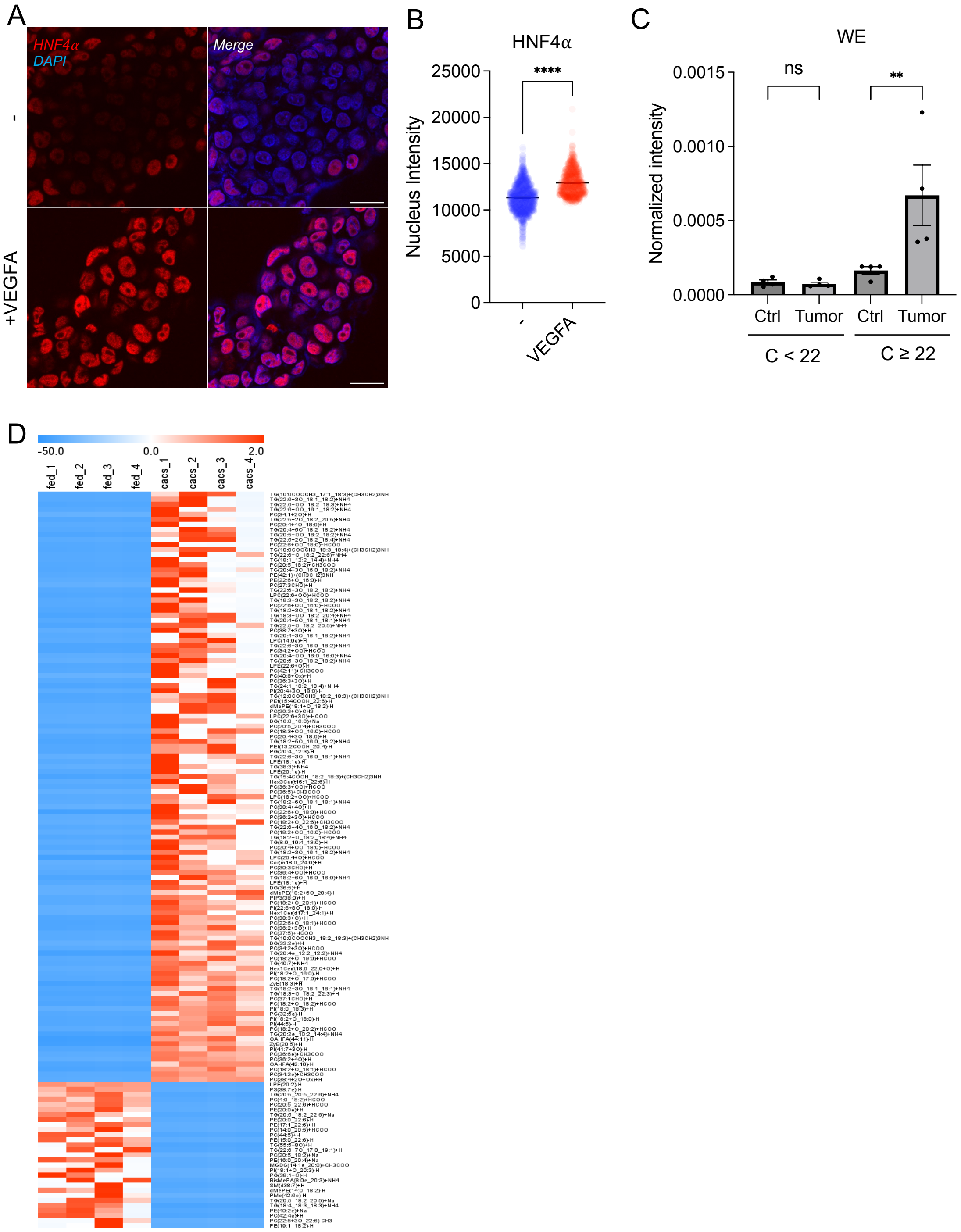


**Figure 7S: VEGF-A regulates lipid metabolism in hepatocytes.**

(A-B) Immunostaining of HepG2 cells treated with or without VEGF-A for 24 hours with Hnf4α and quantification of Hnf4α intensity in the nucleus, blue color indicates DAPI signal, scale bar represents 20μm. (C) The level of WEs with various fatty acid chain length in the serum of tumor bearing or control mice. (D) Heatmap showing the significantly altered lipid species from the serum of control and tumor bearing mice. Data are presented as mean ± s.e.m.; statistical analysis was performed using Student's t-test, and significance analysis of microarrays was used for (D). Ns is $P > 0.05$, *$P < 0.05$, **$P < 0.01$, ****$P < 0.0001$.
